# Modeling Stereospecific Drug Interactions with Beta-Adrenergic Receptors

**DOI:** 10.1101/2023.10.01.560334

**Authors:** John R.D. Dawson, Kevin R. DeMarco, Yanxiao Han, Slava Bekker, Colleen E. Clancy, Vladimir Yarov-Yarovoy, Igor Vorobyov

**Affiliations:** Biophysics Graduate Group, University of California, Davis, CA; Department of Physiology and Membrane Biology, University of California, Davis, CA; Chemistry Department, American River College, Sacramento, CA; Department of Pharmacology, University of California, Davis, CA; Center for Precision Medicine and Date Science, University of California, Davis, CA; Department of Anesthesiology and Pain Medicine, University of California, Davis, CA

**Keywords:** G protein-coupled receptors, beta adrenergic receptors, beta-blockers, Rosetta structural modeling, molecular docking, molecular dynamics simulations

## Abstract

Beta adrenergic receptors (βARs) are G protein-coupled receptors that control processes as varied as heart rhythm and vascular tone by binding agonists such as norepinephrine to induce downstream signaling pathways. Beta blockers antagonize βARs to downregulate their activity, thus reducing heart rate and lowering vascular tone. We developed new Rosetta structural modeling protocol to develop state-specific models of β_1_AR, expressed in cardiac myocytes, as well as β_2_AR, expressed in the smooth muscle cells of vasculature and other tissues, and their atomistic-scale interactions with beta-blockers using RosettaLigand. We identified structural features of drug – receptor interactions, which may account for their receptor conformational state and drug stereospecific preferences. Furthermore, we estimated structural stabilities of our models using atomistic molecular dynamics (MD) simulations. In our recent study we validated our structural models of norepinephrine-bound β_2_AR and its complex with stimulatory G protein via multi-microsecond MD simulations. Thus, here we mostly focused on state-dependent and stereospecific β_1_AR interactions with beta-blocking drugs sotalol and propranolol. We observed expected inactive receptor state preferences and structural stabilities of our models in MD simulations, but neither those simulations nor RosettaLigand docking could clearly distinguish stereospecific preferences of those drugs. This warrants consideration of alternative hypotheses and enhanced sampling MD simulations, which we discussed as well. Nevertheless, our study provides basis for understanding conformational state selectivity and stereospecificity of beta-blockers for βARs, important pharmacological targets, and may be extended to other drug classes and receptor types.

**Graphical abstract:** Norepinephrine (NE) bound active-state beta-1 adrenergic receptor (β1AR) in complex with the stimulatory G protein (Gs) heterotrimer embedded in a lipid bilayer. When expressed at the plasma membrane, the β1AR is oriented such that the ligand binding pocket (*) is accessible to ligands from the extracellular side (Ex.) of the membrane. The Gsα (red), Gβ (blue), and Gγ (yellow) subunits comprise the Gs heterotrimer. Nucleotides GDP or GTP bind Gα at the P-loop (**). *Inset:* Representative image of NE bound within the orthosteric ligand binding pocket.

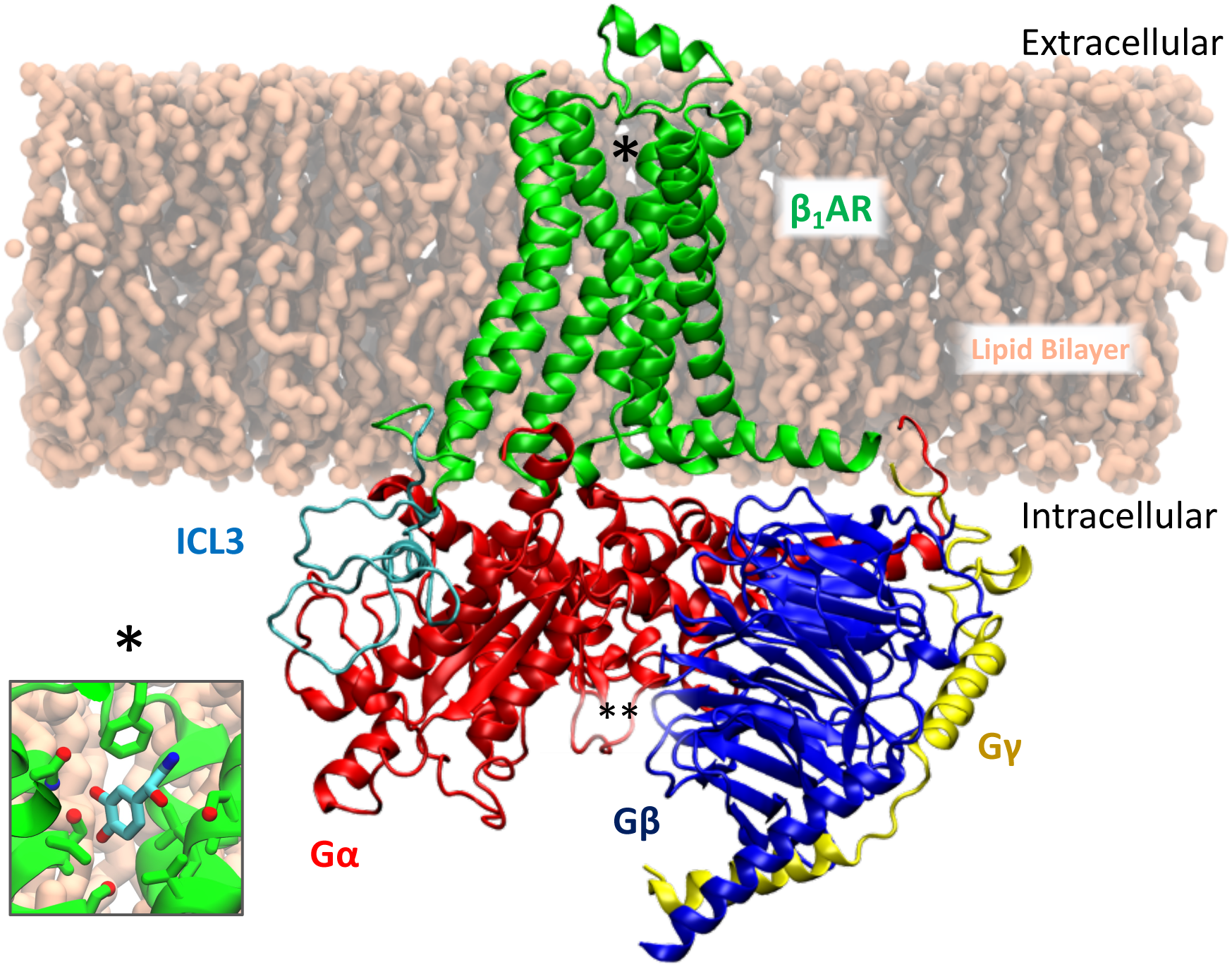

## 1. Introduction

Walter Bradford Cannon, the American physiologist who developed the theory of homeostasis, defined the term “fight-or-flight” to describe the downstream consequences of activation of the Sympathetic Nervous System, or SNS.^1, 2^ SNS stimulation opposes the action of the Parasympathetic Nervous System, or PNS, which mediates the “Rest-and-Digest” and “Breed-and-Feed” processes.^2, 3^ The proper balance of these autonomic systems is fundamental to human physiology and well-being. As society grows more aware of the autonomic nervous system, and as autonomic dysregulation grows more abundant, medicines have been developed to lay a finger on the homeostatic balance between the SNS and PNS.

SNS stimulation of the cardiovascular system increases cardiac output by increasing heart rate, the force of contraction, and conduction rate; consequently, more blood is supplied to the body, a necessary physiological adjustment in circumstances of danger, terror, or exertion.^2^ Excessive stimulation of the sympathetic pathway may induce potentially deadly arrhythmias, especially in situations with underlying cardiac disease.^4^ The discharge of one of either catecholamine neurotransmitters norepinephrine (NE), released from cardiac sympathetic neurons, or epinephrine (Epi), released from the medulla, initiates cardiac sympathetic stimulation.^5^ Alternatively known as “adrenaline” and “noradrenaline,” these substances drive the fight-or-flight response by binding the G-protein coupled receptors (GPCRs) named beta-adrenergic receptors (βARs), which are seven-transmembrane segment proteins found within the cardiac myocytes of the human heart and other vascular tissues.^6^ By binding to beta-adrenergic receptors, NE or Epi induce the receptor to activate the stimulatory G-protein, G_s_. Activation of G_s_ begins the cyliic adenosine monophosphate (cAMP)-induced protein kinase A (PKA) phosphorylation of a multitude of cellular targets that exert electrophysiological changes necessary to increase cardiac contractility.^5^ By inhibiting the binding of NE or Epi to βARs, one prevents the deleterious effects of excessive sympathetic stimulation. Drugs that prevent this action through antagonism are termed beta blockers.^5^ Beta blockers are widely used to treat cardiac irregularities such as atrial fibrillation, myocardial infarction, and heart failure.^5^ There are three βAR subtypes within the human heart: β_1_, which accounts for approximately 75 to 80% of receptors, β_2,_ for 15% to 18%, and β_3_ for only 2% to 3% of βARs.^7^ Though it is critical to note that in the failing human heart, the ratio of β_1_ and β_2_ subtype becomes approximately equal.^8^ Beta blockers have different affinities for each subtype,^9^ and the populations of βARs in non-cardiac vascular tissues differ as well. Namely, β_2_AR is more prevalent in arteries than in the heart and is thus the predominant target of the nonselective hypotensive beta-blocker propranolol ^10^. Beta blockers which predominantly target β_1_AR are “cardio-selective” and are the third generation of beta-blockers.^11^ However, there is another factor that may affect affinity: stereoselectivity.

In our previous study^12^ we discussed the multi-target effects of the antiarrhythmic drug dl-sotalol, a stereoisomeric beta-blocking drug in functional scale cardiac electrophysiological simulations. While both d- and l-sotalol are capable of blocking hERG ion channel current, only l-sotalol is capable of binding beta-adrenergic receptors with a high enough affinity to attenuate sympathetic stimulation at physiological dose.^13^ DeMarco et al. concluded that when modeling hERG potassium channel block, either sotalol stereoisomer is sufficient to induce the markers of deadly arrhythmia in the absence of any simulated sympathetic stimulation.^12^ However, incorporating attenuation of adrenergic signaling into the model eliminates the arrhythmogenic markers for l-sotalol, but not d-sotalol. Indeed, the different selectivity for sotalol has been long known because of the fatal Survival With ORal D-sotalol trial,^14^ wherein the administration of d-sotalol to intentionally block hERG current in patients proved to be deadly. However, the precise molecular mechanism governing this stereoselective block is not known. Furthermore, the prototypical beta-blocker propranolol also exhibits stereospecificity towards beta receptors^15^ and can block hERG channel ^16^, yet whether R-propranolol similarly is also capable of being pro-arrhythmic in the same manner as d-sotalol is unclear. In fact, most beta blockers are administered as racemates. Identifying the molecular mechanisms through which beta-adrenergic receptors exhibit selectivity towards stereoisomers of its ligands via molecular simulations merits consideration.

In this study we discuss the modeling of beta-blockade at the molecular level for the drugs d-sotalol, l-sotalol, R-propranolol, and S-propranolol and receptor interactions with NE to assess the model quality and finally establish whether molecular interactions can be identified, which govern stereoselectivity attributable to cardiac risk (**Table 1**). To do so requires models of the beta-adrenergic receptor subtypes β_1_ and β_2_, which predominate cardiac sympathetic stimulation in healthy and pathological conditions. As it is unclear whether these antagonists prefer the active or inactive state conformations or whether the stereospecific selectivity manifests as such, models of both the active and inactive receptor are necessary for either subtype. It is further important to consider the participation of the stimulatory G-protein (G_s_) in maintaining an active state for drug binding; therefore, active state models with and without the G_s_ heterotrimer are considered. In this study we will focus on sotalol and propranolol interactions with β_1_AR, since in our recent paper we provided detailed molecular simulation analysis of β_2_AR with G_s_ and norepinephrine.^17^ However, the details of structural model development for both β_2_AR and β_1_AR systems, omitted in that study, will be presented here.

**Table 1.**
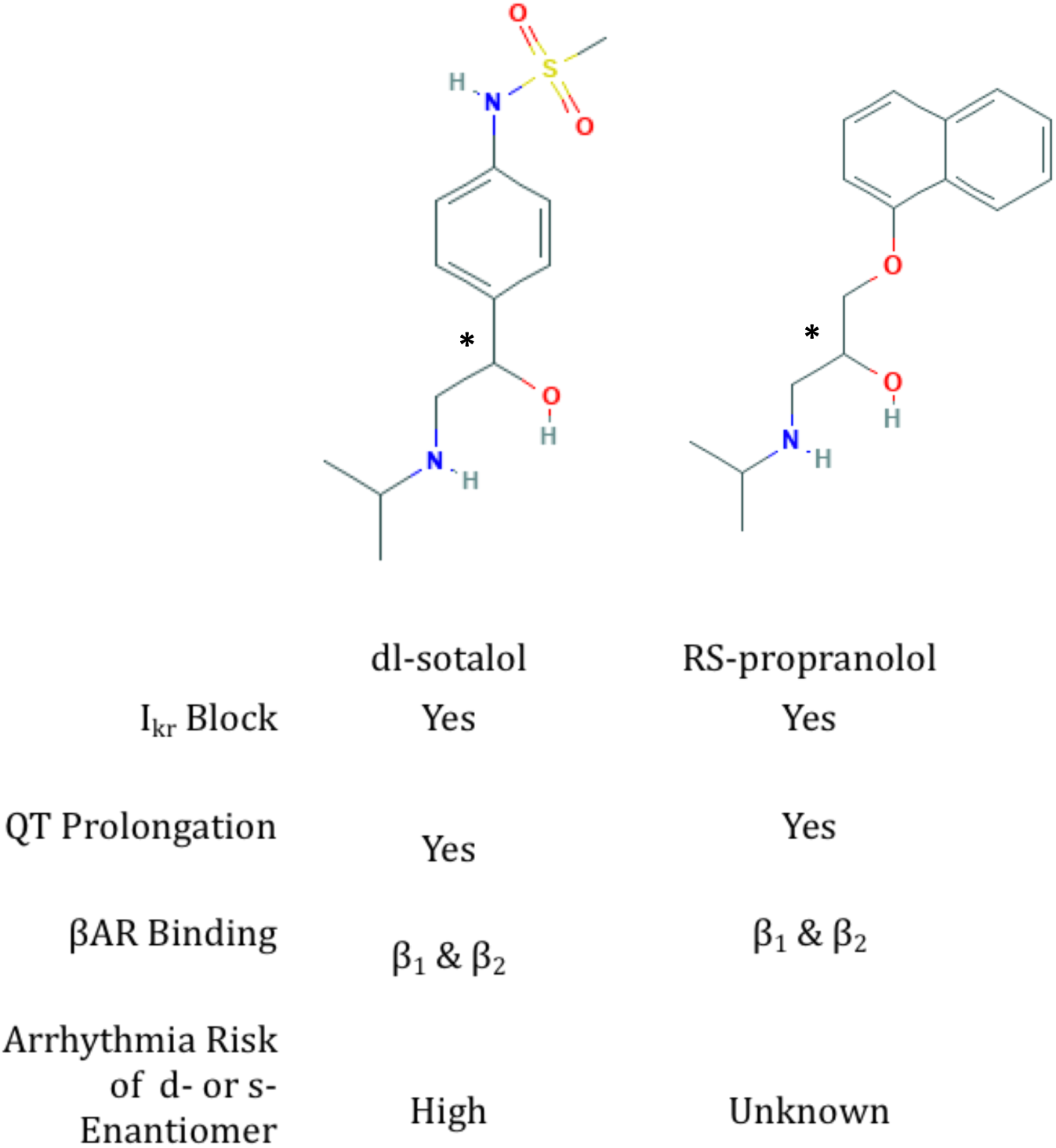
Structure and arrhythmogenic properties of racemic beta blockers dl-sotalol and RS-propranolol. Chiral centers are denoted with asterisk.

|  |  |  |
| --- | --- | --- |
|  | dl-sotalol | RS-propranolol |
| I<sub>kr</sub> Block | Yes | Yes |
| QT Prolongation | Yes | Yes |
| βAR Binding | β<sub>1</sub> & β<sub>2</sub> | β<sub>1</sub> & β<sub>2</sub> |
| Arrhythmia Risk<br>of d- or s-<br>Enantiomer | High | Unknown |

## 2. Methods

### 2.1. Preparation of β_2_AR & β_2_AR-G_s_ templates

Multiple structures of beta-adrenergic receptors and g-proteins are available on the Protein Databank (PDB), though not all of them are derived from human cells. While the sequences of bovine and rat G-protein subunits are identical to those of *Homo sapiens*, the same is not true for the β_1_AR, and only the turkey-derived structures were available at the time this work began. Furthermore, physiologically impactful components such as long and flexible intracellular loop 3 (ICL3) are not resolved in any of them because of their intrinsic disorder. In order to model the unresolved regions of the beta-adrenergic receptor protein, templates must be prepared from available experimental structural templates which may further differ in their originating organism. Therefore, models were prepared identically in preparation for homology modeling to maintain consistent methodology for *de novo* modeling, but only the β_1_AR model is a true homology model, whereas available *Homo sapiens* protein structures were used for β_1_AR models.

The published X-ray crystallographic structure of adrenaline-activated human β_2_AR bound to a high-affinity camelid antibody (PDB: 4LDO) ^18^ was obtained from the PDB to serve as a template for the activated receptor model. For the protein complex model incorporating the G_s_ heterotrimer, its template was isolated from the 3D coordinates of the X-ray crystallographic structure of the β_2_AR-Gs complex bound to agonist P0G (PDB: 3SN6).^19^ The 3D coordinates were obtained as biological assemblies oriented by the Orientations of Proteins in Membranes (OPM) database to be used for molecular dynamics simulation^20^. The adrenaline-bound receptor 4LDO was aligned to 3SN6 using UCSF Chimera^21^ Matchmaker and then used to replace the P0G-bound receptor of 3SN6; then all ligands and non-native protein fragments including the camelid antibody were removed.^21^ The resulting template structure consisted of the beta-adrenergic receptor isolated from PDB ID 4LDO in complex with the G_s_ heterotrimer from PDB ID 3SN6. This complex was then assessed for steric clashes within van der Waals radii and was found free of collisions and thus suitable for structural modeling.

### 2.2. β_2_AR & β_2_AR-G_s_ loop rebuilding

The β_2_AR structure 4LDO has unresolved ILC3 as well as the termini and similarly disordered portions of the G-protein. These regions were modeled *de novo* using the ROSETTA implementation of fragment-based cyclic coordinate descent method (CCD)^22^. Target sequences for remodeled regions of either the human β_2_AR (**Fig. S1**) and the G_s_ heterotrimer (**Fig. S2**) were obtained from UniProt^23^ and visualized using Jalview^24^. The Hybridize Mover of Rosetta comparative modeling (RosettaCM) was applied with the Rosetta Membrane Energy Function^25^ to generate 10,000 decoy models of loop-rebuilt β_2_Ar-G_s_ from which a candidate model was selected for further simulation.^26, 27^ Rosetta clustering analysis with root-mean-squared deviation values was used to assess the convergence of models into different microstates using a radius of 2.5 Ak^28^. The lowest-energy model of the most populated cluster was selected as a candidate model for additional refinement (**Fig. S3A&B**).

Energy minimization was then applied to relax the backbone and sidechain region of rebuilt portions of β_2_AR-G_s_ complex. One thousand energy-minimized decoys were generated from the sequence-complete β_2_AR-G_s_ using the Rosetta FastRelax application with the membrane energy function^29^. Relaxation was permitted only to residues that were modeled *de novo* or residues within 3.5 Ak of the α5 helix of G_s_α subunit of the G_s_ heterotrimer. The lowest energy decoy was then selected for ligand docking and MD simulations.

### 2.3. Preparation of active β_1_AR-G_s_ and inactive β_1_AR templates

The published X-ray crystallographic structure of isoprenaline-activated, turkey β_1_AR bound to nanobody Nb80 (PDB: 6H7J) was obtained from the PDB as biological assemblies lipid membrane oriented by the OPM database.^30^ To form the human β_1_AR-G_s_ complex, the sequence-complete G_s_ heterotrimer model was isolated from our β_2_AR-G_s_ complex model derived from its X-ray crystallographic structure (PDB: 3SN6), ensuring continuity of the G-protein conformation between models^19^. The isoprenaline-bound receptor from PDB ID 6H7J structure was aligned to our β_2_AR-Gs candidate model using UCSF Chimera^21^ Matchmaker to ensure consistency in orientation with the G-protein-free template and to replace the β_2_AR receptor model with the activated turkey β_1_AR to create a β_1_AR-G_s_ templates.^21^ All ligands and non-native proteins including isoprenaline and nanobody Nb80 were then removed. The resulting template structures consisted of the turkey beta_1_-adrenergic receptor 6H7J in complex with our previously modeled G_s_heterotrimer. The turkey β_1_AR-G_s_ complex was assessed similarly for steric clashes using van der Waals radii before homology modeling.

### 2.4. β_1_AR & β_1_AR-G_s_ homology modeling

Homology modeling was used to generate putative receptor models of human active-state β_1_AR and β_1_AR-G_s_ complex from their homologous turkey-derived template structures using a target sequence for human β_1_AR obtained from Uniprot. Sequence similarity between the human and turkey structures was (48.65%), permitting effective use of comparative modeling via RosettaCM. Much of the divergence between the β_2_AR and β_1_AR sequences originate within the length of the ICL3; β_1_AR 6H7J structure lacks around sixty residues in the ICL3 region, compared to twenty-seven in β_2_AR 4LDO. Therefore, multiple sequence alignment using Clustal Omega was first performed to determine the identical start and end points for structural modeling.^31^ Human β_1_AR was modeled to complete the ICL3 and to match with the N- and C-termini of our β_2_AR model, which has the identical sequences in those regions. Partial threading was first performed to generate a sequence-correct human template model fitted into the geometry of the turkey structure and in accordance with the sequence alignment. Then RosettaCM was applied: fragment-based method for protein structure building using templates sampled from a pre-generated fragment library to complete missing regions in a three-stage protocol that completes and refines the modeled geometry in a Monte Carlo trajectory. First, fragment recombination is used to construct a sequence-complete template. Recombination is performed by inserting fragments in torsion space from randomly those randomly selected from the fragment library, and then scored using the low-resolution centroid scoring function. Second, to optimize this geometry and correct for unrealistic backbone lengths, de novo fragments or library-based fragments are randomly super-imposed to substitute particularly distorted regions of the first stage output and then concludes with another round of full-backbone energy minimization using a cartesian space centroid energy function. After 1,000 runs, the lowest energy structure is passed to the third stage. In the third stage, side chains are incorporated into the model using Rosetta’s Monte Carlo combinatorial side-chain optimization, and subsequently minimized using “FastRelax.” The Rosetta Membrane Energy Function REF15^32^ was used for centroid and full-atom scoring.^27^ 10,000 decoy models of β_1_AR and β_1_AR-G_s_ were generated from which candidate models were selected ^26, 27^. Rosetta clustering analysis was used to assess the convergence of models into different microstates based on their RMSDs using a radius of

2.5 Ak.^28^ The lowest-energy decoy of the most populated cluster was selected as the candidate model for additional refinement using Rosetta FastRelax. ^29^ 1,000 energy-minimized decoys were generated for either candidate homology model. Relaxation was permitted only to residues that were modeled *de novo* or residues within 3.5 Ak of the α5 helix of G_s_α subunit of the G_s_ heterotrimer. The lowest energy decoy was then selected for ligand docking and MD simulations.

For the inactivated state, the isoprenaline-bound turkey β_1_AR structure (PDB: 2Y03) “with stabilizing mutations” was chosen as a template and prepared identically, but without the G_s_ heterotrimer.^33^ Though isoprenaline is an agonist, this structure adopts an inactivate-like conformation, and because it lacks any auxiliary proteins such as nanobodies within the cytosolic region it may serve as a control for assessing allosteric effects in subsequent molecular dynamics simulations.

### 2.5. Preparation of Human β_1_AR & β_!_AR-Gs Templates from Crystallographic Structures

The release of a more-contemporary membrane energy function “Franklin2019” ^32^, and critically important the publishing of multiple structures of the human β_1_AR, necessitated a second pass at the modeling both states of β_1_AR, and their interactions with the G-protein. Xu et al. published crystal structures of the human β_1_AR in complex with epinephrine (PDB: 7BU6) and the antagonist carazolol (PDB: 7BVQ) bound to the nanobody 6B9.^34^ These structures were selected as templates for active and inactive state models of β_1_AR, respectively. The previously modeled G_s_ heterotrimer derived from PDB structure 3SN6 was used to form a human β_1_AR-G_s_ complex template. All model coordinates were obtained as biological assemblies oriented by the Orientations of Proteins in Membranes (OPM) database, but were subsequently aligned to the previously developed β_2_AR-G_s_ model and cleaned of ligands and all non-native peptides using UCSF Chimera^21^ to ensure consistent orientation.^20, 21^. The human β_1_AR-G_s_ complex was assessed for steric clashes using van der Waals radii before proceeding to loop modeling.

The β_1_AR structures lack coordinates for the ICL3, which in β_1_AR is substantially longer and potentially more disordered than its β_2_AR counterpart. To harness the new membrane energy function “Franklin2019”, a stepwise protocol was devised in place of comparative modeling to achieve better convergence. To model the sixty-residue long ICL3 and eleven residues of C-terminus *de novo*, loop reconstruction was performed using kinematic closure with fragments in staged process.^35^ 1,000 decoys of the ICL3-added receptor models were generated and clustered using RMSDs with a radius of 5 Ak, and a candidate model was selected from the lowest energy cluster. 4,000 decoy models with added C-terminus were then generated, and a top candidate model was selected from the lowest energy cluster after clustering using RMSDs with a radius of 0.8 Ak. Relaxation was performed using FastRelax and the Franklin2019 membrane energy function after ligand docking was performed as described below.

### 2.6. RosettaLigand docking of endogenous norepinephrine and beta blockers to βAR models

RosettaLigand^36^ was used to simulate the docking of ligands to β-adrenergic receptors. Up to 200 rotamers and Rosetta energy function parameters were generated for the ligands norepinephrine, R-propranolol, S-propranolol, d-sotalol, and l-sotalol by OpenEye Omega ^37^ and ROSETTA scripts. A box size of 5 Ak was used for ligand transformations such as rotation, translation, and conformational changes along with a 7 Ak ligand distance cutoff for side chain and backbone reorientations (with <0.3 Ak C_α_ restraint). 50,000 docked poses were generated in each run with the top 10% selected by total score, out of which the fifty lowest-interfacial score decoys were verified for their convergence and accuracy with the crystallized ligand of their original PDB template structure. Subsequent molecular dynamics simulations were conducted using the absolute lowest-interfacial score structure to serve as the candidate model, unless otherwise specified. Analysis of binding sites was performed in UCSF Chimera^21^ and using the Protein-Ligand Interaction Profiler or PLIP^38^.

### 2.7. Molecular dynamics simulations

Molecular dynamics were used to better understand state-specific dynamics and stability of the human beta-adrenergic receptor complexes and the nature of their interaction with beta blockers. CHARMM-GUI Membrane Builder^39, 40^, an online web service for establishing initial systems for MD simulations using the CHARRM force field^40^, was used to create systems of hundreds of thousands of atoms. The CHARMM-GUI Membrane Builder follows a five-step protocol where PDB coordinates are first read into the web server and then oriented according to parallel *XY*-planes representing the upper and lower lipid bilayer leaflets. The dimensions of the system are then determined, including the minimal extent of water needed. Fourth, the individual components are calculated and built separately: the lipid bilayer is placed around the protein, water molecules are placed to solvate the protein, and ions are placed using Monte Carlo sampling to populate the solvent according to a prescribed concentration. Fifth, these individually built components are assembled into a single system. Sixth, the system and pre-determined equilibration protocol are provided to the user, though this equilibration protocol is inadequate, and the presented systems underwent additional equilibration.

Using this methodology, each beta-adrenergic receptor model was embedded in a heterogenous lipid bilayer of palmitoyl-oleoyl-phosphatidylcholine (POPC) and palmitoyl-oleoyl-phosphatidylserine (POPS) of approximately two hundred lipids per leaflet. This composition was adopted from previous simulations by Dror et al.. 2015^41^, and consisted of an upper leaflet of POPC lipids and a bottom leaflet 70:30 mixture of POPC:POPS lipids. Protonation states, terminal group patching, histidine protonation, and lipidations were similarly derived from Dror et al., 2015.^41^ Critically, each receptor was protonated at the equivalent residues Glu147 and Asp155 for β_1_AR, or Glu122 and Asp130 for β_2_AR. S-palmitoylation is specified Cys392 in β_1_AR and Cys341 in β_2_AR. For all simulations incorporating the G-protein heterotrimer, the three additional residues were further lipidated: Gly2 of Gsα was myristoylated and Cys3 palmitoylated, while Cys68 of Gγ was geranlygeranylated. Beta-adrenergic receptors were given acetylated N-terminal and methylamidated C-terminal patches. When included, the heterotrimeric G-protein termini were also patched. Gsα was given a standard carboxylic C-terminus, but as the N-terminus of G_s_α forms the crucial α-5 helical insertion that mediates receptor association, G_s_α was given neutral acetylated N-terminal patch. Gβ had also an acetylated N-terminus and a standard carboxylic C-terminus. Gγ had an acetylated N-terminus and methylated C-terminus as this region inserts into the lower leaflet. All histidine residues were epsilon-N protonated, except residues 225 and 331 of Gβ which were instead delta-N protonated. The systems were solvated with 150 mM aqueous NaCl using the CHARMM36m all-atom protein force field. ^42^ C36 lipid force field, ^43^ standard CHARMM ion parameters with latest non-bonded fix (NBFIX) terms^44^ and TIP3P water model^45^ were used as well. Parameters for norepinephrine and RS-propranolol were first obtained using the general CHARMM force field (CGENFF) ^46^ for small molecules via CGENFF program.^47^ Cationic norepinephrine parameters were further optimized using the force field toolkit (FFTK) plugin^48^ for Visual Molecular Dynamics (VMD) software^49^ using Gaussian software^50^ for reference quantum mechanical calculations. CHARMM compatible force field parameters for cationic and neutral sotalol were generated by Drs. Kevin DeMarco and Igor Vorobyov and were previously published. ^51^ MD simulations were performed with Nanoscale Molecular Dynamics (NAMD) software^52^. MD simulations were conducted in the *NPT* ensemble at 1 atm pressure and 310 K. Nose-Hoover extended system method ^53^ was used for temperature control, whereas the Langevin piston method^54^ was used for pressure control. Particle mesh Ewald (PME) for long-range electrostatic interactions^55^ along with standard non-bonded cutoffs recommended for lipid membrane containing systems^43^ were used as well. SHAKE algorithm was used to constraints bond containing hydrogen atoms ^56^ allowing to use a 2 fs time step.

All MD simulation systems underwent a 40-nanosecond-long staged equilibration protocol with gradually reduced restrains as shown in **Table S1** to allow time for *de novo* modeled regions of the proteins to rearrange, without risking the stability of the protein core, and to establish reference data for the drug binding poses. To extend MD simulation time to the order of microseconds, equilibrated systems were run on the Anton 2 supercomputer^57^. Visualizations and analysis were performed in VMD^49^ and using in-house scripts. The details of MD simulations of norepinephrine-bound β_2_AR and β_2_AR-G_s_ systems are provided in our previous study.^17^, while only d- and l-sotalol-bound β_1_AR systems are featured here.

## 3. Results

### 3.1 Modeling active and inactive state beta adrenergic receptors using RosettaCM

Ten-thousand decoy models were generated for both active and inactive states of human β_2_AR and β_1_AR structural models derived using human β_2_AR or turkey β_1_AR template structures with RosettaCM (see **Figure 1**). For active-state models, bound stimulatory G (G_s_) protein was present based on its structure from the β_2_AR-G_s_ complex (PDB ID: 3SN6). The top four most-populated clusters were examined for each of them, but no clear relationship between root-mean-square deviation (RMSD) from the top-scoring decoy and a given decoy’s score could be established (**Fig. S3 B&D**).

**Figure 1.**
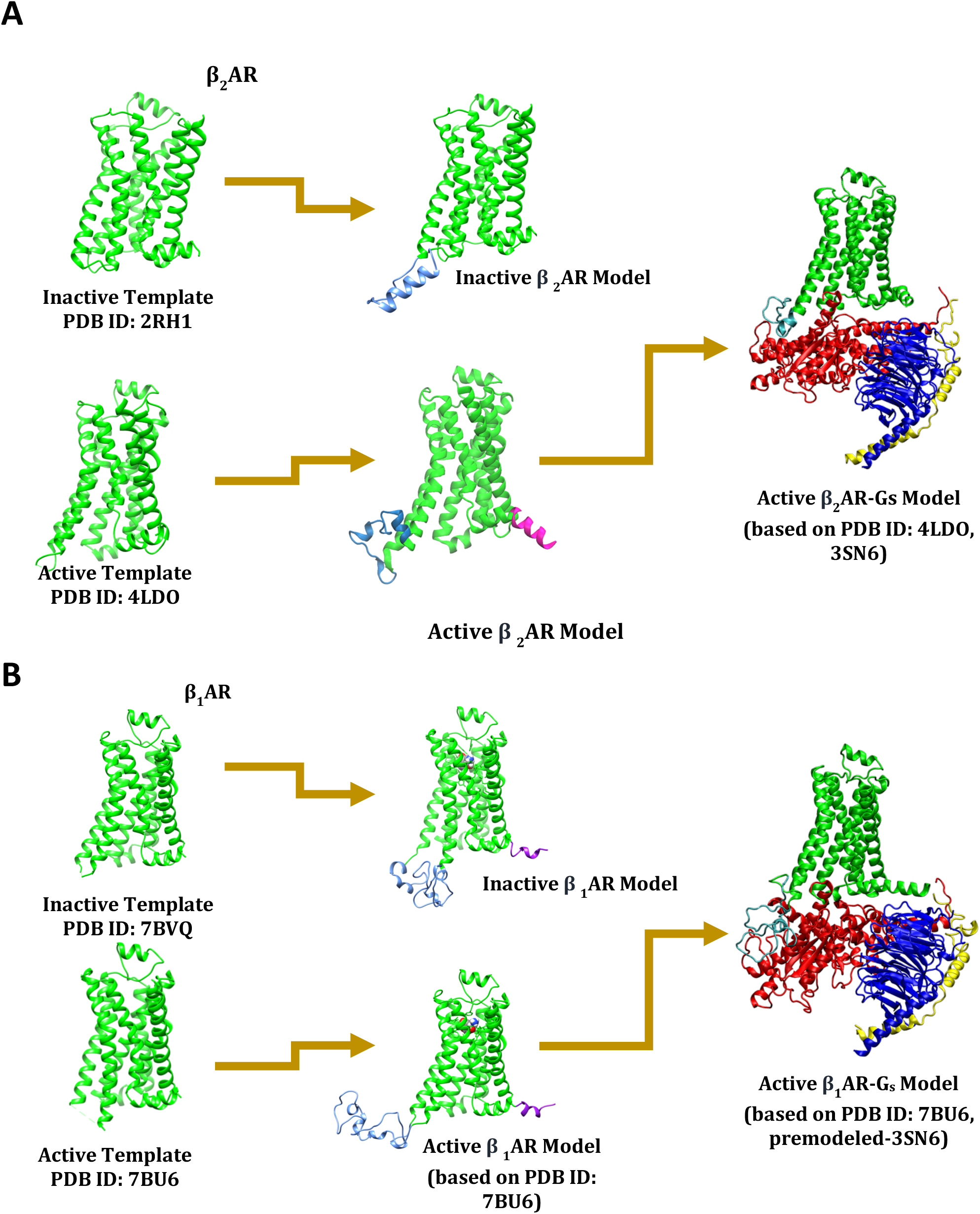
Schematics of modeling protocol for (A) β_2_AR and (B) β_1_AR and their complexes with the stimulatory G (G_s_) protein developed Using RosettaCM. Template models are prepared from published crystal structures for respective states. For active state models loop modeling and FastRelax are performed in the presence of the stimulatory G protein.

In the case of active state β_1_AR, the intracellular loop 3 (ICL3) was so disordered such that clusters were entirely governed by the orientation and direction of ICL3 and not its secondary structure. Therefore, the candidate model was chosen based on the assumption that the ICL3 would not readily penetrate region, which Rosetta treats as implicit membrane. The candidate model for β_1_AR was thus the top scoring decoy from cluster 2 (**Fig.S3 A**). Curiously, loop remodeling with the membrane energy function permitted loops to traverse into the implicit membrane. As G_s_α was present during loop remodeling, the ICL3 could not similarly be built downwards into solvent as it could be outward or upward.

As the ICL3 of β_2_AR is significantly shorter than that of β_1_AR (27 residues vs. ∼60 residues in β_1_), the loop would not traverse into an implicit membrane region, and clusters therefore aggregated either away from or adjacent to G_s_α. (**Fig. S3 C**). In this instance, the top scoring decoy of the lowest-energy cluster was selected as the candidate model (**Fig. S3 C**).

The same protocol and selection criteria were applied to select an inactive state β_2_AR model (**Fig. S4, top**) in the absence of the G_s_ heterotrimer. For inactive β_1_AR the lack of the G-protein meant *de novo* modeling of an entirely unrestrained 60-residue long ICL3. Clustering yielded twenty low-population clusters (**Fig. S4 B**) that lacked any predominant secondary structure aside from the top-scoring model (**Fig. S4 C**) which was selected to be the candidate model and relaxed.

### 3.2 Docking of norepinephrine into RosettaCM-derived active and inactive state beta adrenergic receptors

To validate preservation of the ligand binding pocket and to generate representative models of functional receptor complex with its endogenous ligand, neutral and cationic forms of norepinephrine (NE), were docked using ROSETTA-Ligand^36^ into active-state models of both receptors as well as the inactive state model of β_2_AR **(Fig S5)**. Top fifty poses for all conditions except neutral norepinephrine in active β_2_AR exhibited tight binding. Only in the case of cationic NE docking to active β_2_AR was the crystallographic epinephrine binding pose (**Fig S5 A**) consistently recapitulated. The order of peak probabilities of interfacial scores or interaction energies (IE) would suggest that the more similar the crystal ligand is to the docked ligand, the lower (more favorable) is the IE. The mean IE of docked norepinephrine into a formerly epinephrine occupied binding pocket from PDB ID 4LDO is more favorable than docking into a formerly isoprenaline (agonist) or propranolol (antagonist) bound models based on PDB IDs 6H7J and 6PS5, respectively^30, 58^. However, their probability distributions nearly overlap (**Figure S5 D**). This trend is abolished for the case of neutral norepinephrine, despite mostly tight geometric convergence within the binding pocket, a change likely attributable to expected much higher affinity of cationic NE binding to β_2_AR, which cannot be directly assessed using Rosetta energy scoring function.

### 3.3 Docking of stereoisomeric beta blockers into RosettaCM-derived inactive-state beta adrenergic receptors

The top fifty scoring poses of docked d- and l-sotalol docked to inactive β_1_AR occupy nearly identical regions of the binding pocket and their IEs in Rosetta Energy Units (REU) are statistically indistinguishable: l-sotalol: −10.64 +/- 0.71 REU versus d-sotalol −10.67 +/- 0.37 REU, averaged over the top 50 poses (**Fig. 2A**). Their top binding poses both have their sulfonamide moiety in the same orientation within the binding pocket, and coordinating with Ser228 but not Ser232. Both serine residues, denoted Ser S^5.24^ S^5.46^ Ballesteros-Weinstein nomenclature^59^ govern catechol hydroxyl recognition^60^ as seen in the crystal structures for β_1_AR-Epinephrine(PDB ID: 7BTS) and β_1_AR-Norpinephrine(PDB ID: 7BU6), and the and crystal structures of β_2_AR-Epinephrine(PDB ID: 4lDO) (**Fig. 2 A&B**)^34^.

**Figure 2.**
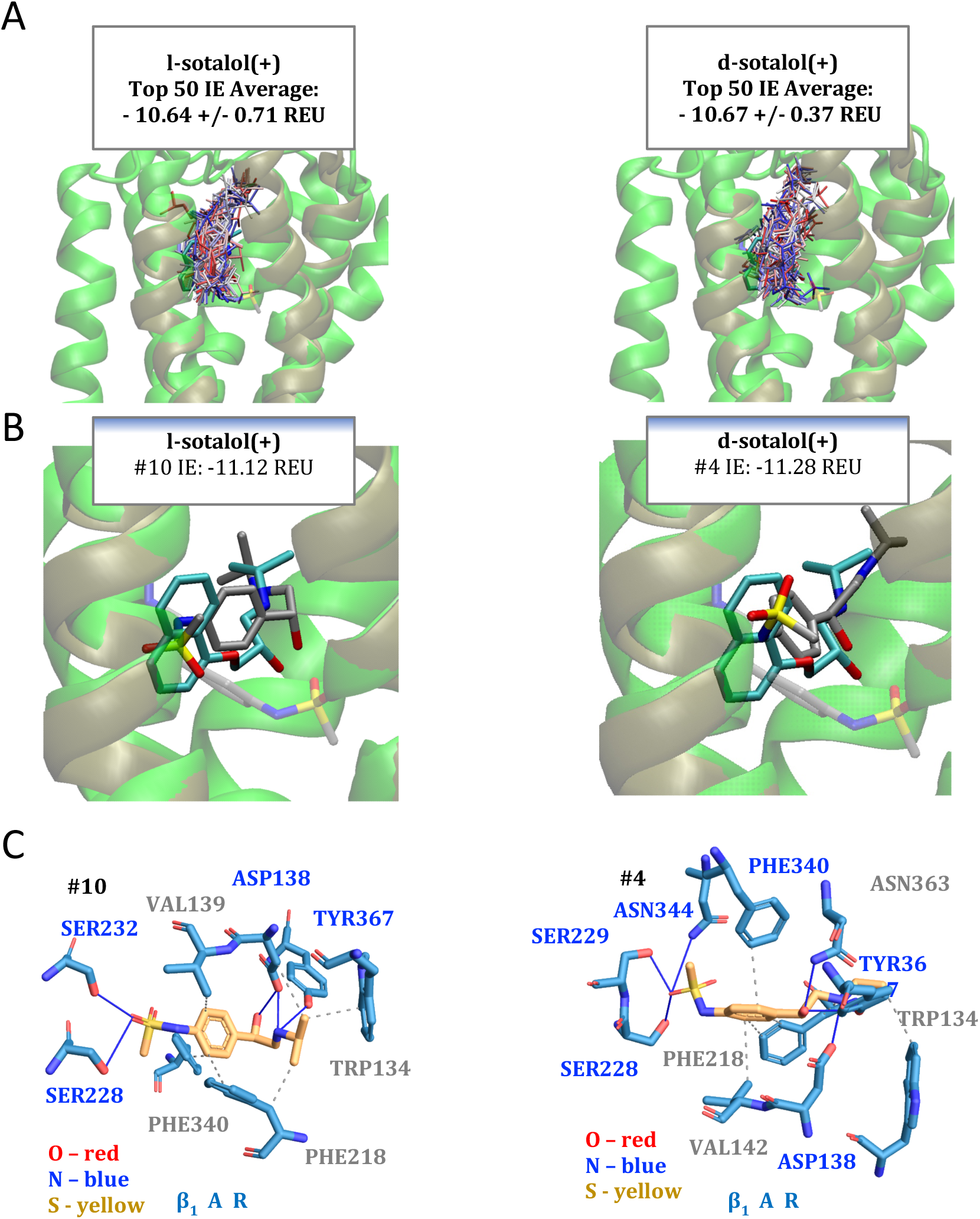
Docking of dl-sotalol stereoisomers into inactive human β _1_AR homology model. **(A)** Top fifty poses of l-sotalol (left) and d-sotalol (right) colored from red to blue as first to fiftieth lowest energy. β_2_AR crystal structure from PDB ID 6PS5 (tan) with bound cationic S-propranolol (cyan) is shown as a reference in panels A and B. **(B)** β_1_AR protein model in green with tenth top-scoring pose of l-sotalol (left, gray) or forth scoring pose of d-sotalol (left, gray) and their respective initial positions (transparent). **(C)** PLIP analysis showing interacting residues of tenth scoring pose of l-sotalol (left) and fourth scoring pose of d-sotalol (right). Nonpolar interactions are denoted by gray dashed lines, and hydrogen-bonding interactions by blue lines.

Cationic RS-propranolol, being a larger molecule, occupied a larger volume of the binding pocket, including a region deeper into β_1_AR interior, which is seldomly sampled by sotalol (**Fig. 3A&B**). The mean IEs of the top fifty poses for both stereoisomers lie within error of one another (S-propranolol: −13.19 +/- 0.34 REU versus d-sotalol −13.31 +/- 0.64 REU). Both sets of docking data tougher indicate that IEs computed using RosettaLigand may not account for experimentally known stereospecificity against the β_1_AR homology model. Therefore, molecular dynamics simulations were conducted based upon the candidate sotalol-β_1_AR complexes shown in **Fig. 2B**.

**Figure 3.**
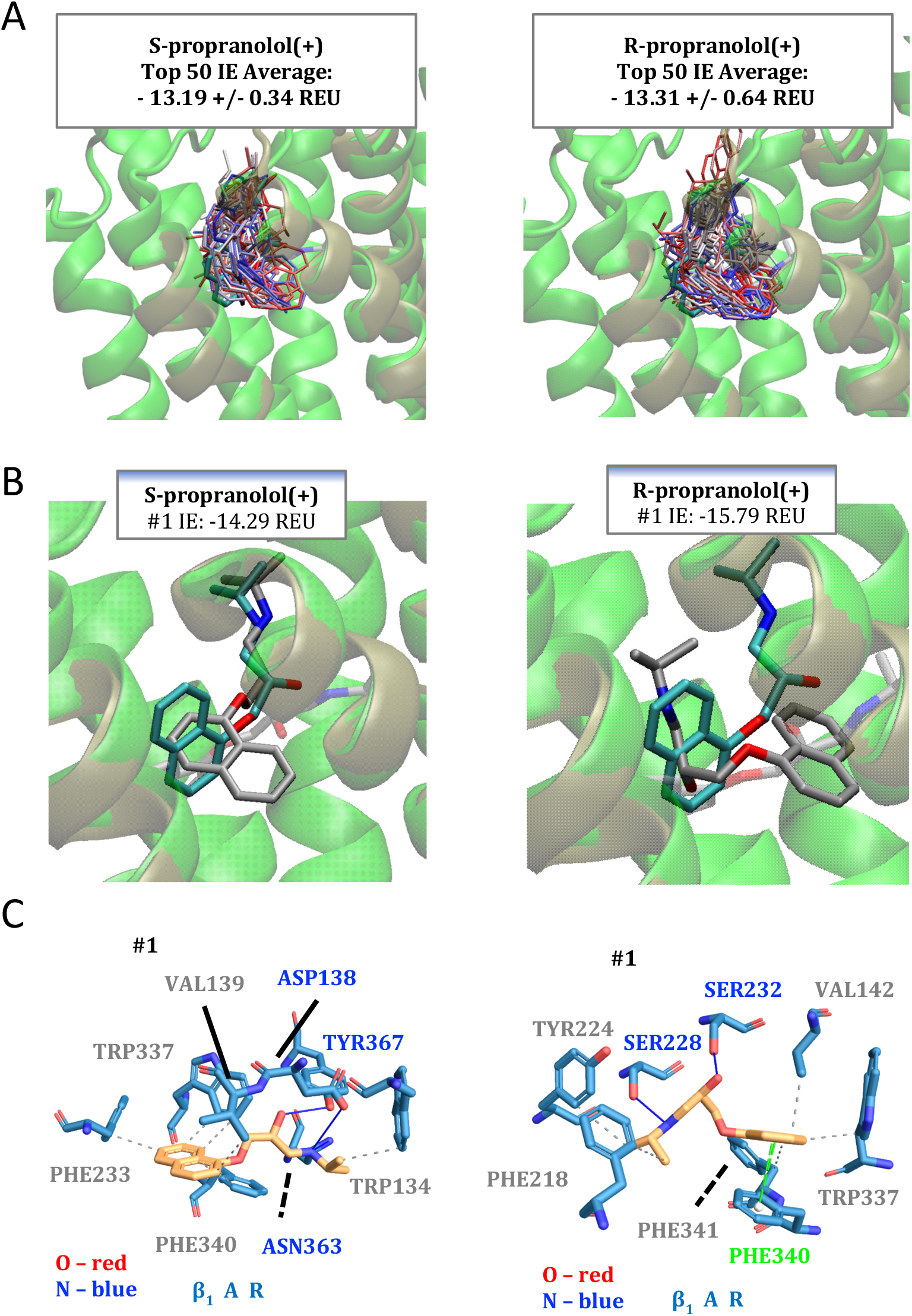
Docking of RS-propranolol stereoisomers into inactive human β_1_AR homology model. **(A)** Top fifty poses of S-propranolol (left) and R-propranolol (right) colored from red to blue as first to fiftieth lowest energy. Inactive β_2_AR crystal structure from PDB ID 6PS5 (yellow) with bound cationic S-propranolol (cyan) is shown as a reference in panels A and B. **(B)** β_1_AR protein model in green with top scoring pose of propranolol (solid gray) and initial position (transparent gray). **(C)** PLIP analysis of interacting residues of top poses for S-propranolol (left) and R-propranolol (right). π-stacking interaction is denoted by green line, nonpolar interactions are denoted by gray dashed lines, and hydrogen-bonding interactions by blue lines.

### 3.3 Molecular dynamics of sotalol interactions with the human β_1_AR homology model

To test whether incorporating dynamics into these systems could potentially reveal the stereospecific drug binding preferences, the candidate poses of human β_1_AR homology model docked to either d- or 1-sotalol were simulated using fully atomistic molecular dynamics. However, as the docking simulations suggested, no stereospecific preferences were observed in MD simulations either. **Fig. 3** depicts results of multi-microsecond unbiased MD simulations of either sotalol stereoisomer bound to the inactive-state β_1_AR model. Between the two stereoisomers, l-sotalol reorients most significantly during the simulation, yet both maintain the same orientation within the binding pocket and end the simulation at approximately 4 Ak RMSD with respect to the initial binding pose. Curiously, while both drugs begin with similar positions of their sulfonamide group with respect to the receptor, in the case of d-sotalol this moiety digs deeper into the protein interior and holds a consistent orientation for longer. However, the dynamical consequences of sotalol binding on the receptor structure do not significantly differ between stereoisomers. In both conditions the receptor adopts a more inactivated orientation, with TM6 shifting more inward. With no clear obvious distinction suggesting preferential β_1_AR interactions with l-sotalol over d-sotalol, enhanced sampling MD simulation techniques beyond the scope of this study and/or revision of the structural receptor models are necessary.

### 3.5 Structural modeling of active and inactive state of beta-1 adrenergic receptors

Recently human beta-1 adrenergic receptor structures were published^34^, which eliminates homology modeling as a potential variable in experimental design. Therefore, we developed new models of β_1_AR using those structures as templates and adjusting our previous Rosetta protocol to achieve better convergence and potentially accuracy as well. The protocol for creating models was similar to that in the RosettaCM process described above and is depicted in **Figure 4A**, however *de novo* structural modeling was broken up into two steps of kinematic loop remodeling with fragments. Whereas using our previous approach RosettaCM yielded one model with the ICL3 and the C-terminus rebuilt, this new protocol performed the *de novo* modeling in two separate steps. This permits the best practice of selecting for the most-convergent regions per-segment; clustering is otherwise less effective when assessed by RMSD across the entire protein as was done previously. Furthermore, eliminating RosettaCM meant that template sidechains were unchanged, better preserving the binding site conformation. Lastly, the alternative loop modeling protocol, when applied using the Franklin2019 Rosetta membrane energy function, also eliminated membrane-penetrating loop rebuilds which would otherwise be discarded and worsened sampling. The final active and inactive state protein models, shown docked with norepinephrine in **Fig. 5B&C** (right-hand side) have considerably different ICL3 when compared to those in **Fig. 1B and Fig. S4 D**.

**Figure 4.**
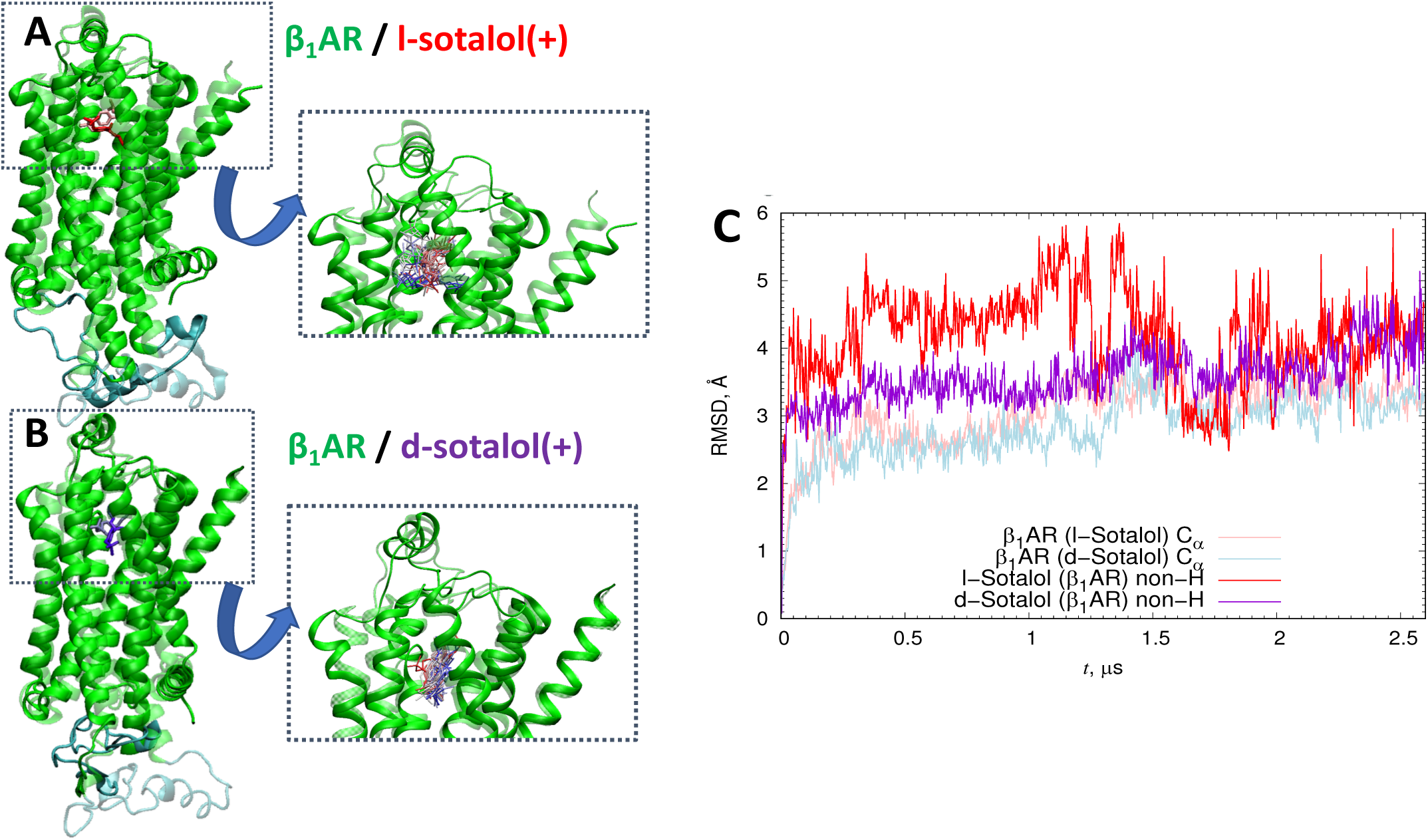
Multi-microsecond-long all-atom MD simulations of cationic d- and l-sotalol bound to inactive beta-1 adrenergic receptor homology model. Initial (transparent) and final (solid) positions of the receptor and drugs (**A**) cationic l-sotalol (red) and (**B**) d-sotalol (blue). Inset images are timeseries of individual poses of either drug taken from the 2.5-microsecond-long simulations and are colored by from the beginning of the trajectory (red) to the final frame (blue). (**C**) Root-mean-square deviation (RMSD) of protein C_α_ atoms as well as either d- or l-sotalol non-hydrogen atoms from their initial docked positions with respect to the receptor.

**Figure 5.**
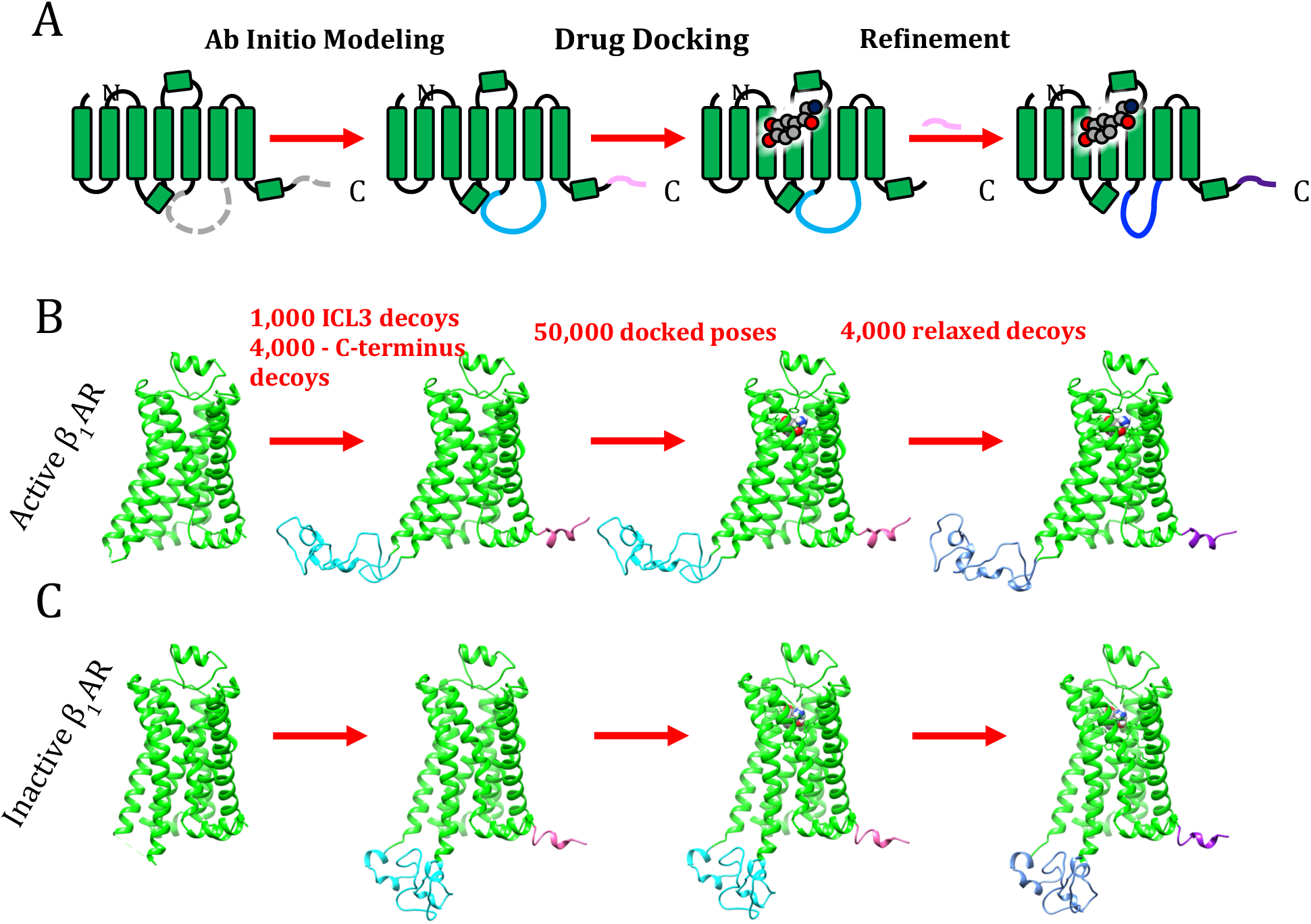
Loop modeling protocol using human β_1_AR structural templates and Franklin2019 Rosetta membrane energy function. (**A**) This protocol uses kinematic loop closure to generate putative structures of the ICL3 and C-terminus of β_1_AR and was applied for both (**B**) active and (**C**) inactive states of human β_1_AR.

To validate stability of the model, the active receptor was simulated in molecular dynamics without the G-protein after being docked with cationic norepinephrine in accordance with the previous RosettaLigand and MD simulation protocols (**Fig. S6**). The final frame of the MD simulation at ∼92 ns illustrates a loss of some secondary structure of the ICl3 as it extends in the aqueous solution. Norepinephrine binding remained intact, and so we proceeded to docking beta blockers to our new human β_1_AR structural models.

### 3.6 Docking of stereoisomeric beta blockers into active and inactive state beta-1 adrenergic receptor models

Beta blockers RS-propranolol and dl-sotalol were docked into the new human β_1_AR structural model using a RosettaLigand protocol outlined above. When docked to the active-state β_1_AR model, the same findings as determined via the receptor homology model held: the most favorable interfacial scores between d- and l-stereoisomers of sotalol were indistinguishable (see **Fig. 6A**).

**Figure 6.**
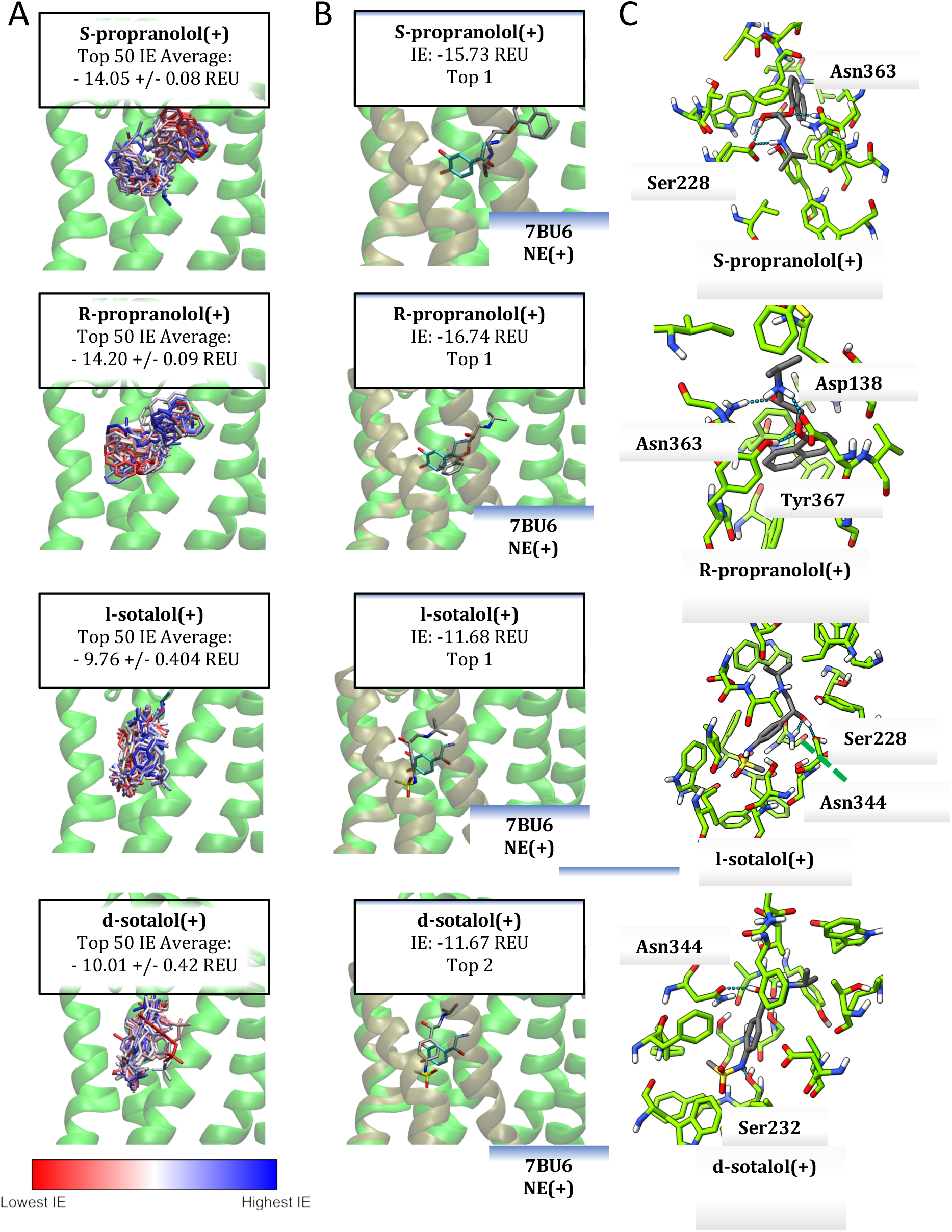
RosettaLigand docking stereoisomers of sotalol and propranolol into active-state human beta-1 adrenergic receptor. (A) Top fifty poses of cationic drug docking into the active human β_1_AR model colored from lowest interaction energy, IE (red, most favorable) to highest IE (blue, least favorable). (**B**) Candidate pose selected from top ten scoring poses compared to crystalline norepinephrine (NE) (cyan) as bound to the original template model (tan) from PDB ID:7BU6. (**C**) Contact PLIP analysis of the candidate pose for each drug with interacting residues labeled and H-bonds depicted by dotted blue lines.

When examining the average interfacial energy of the top fifty poses, we found that R- and S-propranolol had more favorable interfacial scores (R-propranolol: −14.05 +/- 0.08 REU, S-propranolol: −14.20 +/- 0.09 REU.) than sotalol (l-sotalol: −9.76 +/- 0,40 REU, d-sotalol: − 10.01 +/- −.42 REU), although the direct comparison might not be possible due to use of knowledge-based terms in the Rosetta energy function. Though top 1 pose of d-sotalol scored higher than the top 1 of l-sotalol (−13.46 REU, not shown), the top 2nd pose of d-sotalol was selected as a candidate model on account of orientation. For all candidate poses, binding orientation is distinct from co-crystalized NE from the original template structure (**Fig. 6B**). One of either opposing asparagine residues Asn344 and Asn363, associated with receptor-type selectivity, and one of either serine residues Ser228 or Ser232, associated with catechol hydroxy recognition, are forming hydrogen bonds with the ligand for all beta blockers(**Figure 6C)** ^60^.

Average interfacial scores determined from the top fifty poses against the inactive state β_1_AR model indicate that RS-propranolol binding worsened, while l-sotalol binding improves, and d-sotalol remains constant (see average IEs in **Figs. 6A and 7A**). Regarding differences between stereoisomers, for S- and R-propranolol it lies within an uncertainty (S-propranolol: −13.97 +/- 0.88 REU versus R-propranolol: −12.68 +/- 0.53 REU) (**Fig. 7A).** The average interfacial scores for sotalol also overlap, suggesting no significant difference (l-sotalol: −10.87 +/- 0.098 REU versus d-sotalol: −10.18 +/- 0.63 REU) (**Fig. 7A**). The top pose for S-propranolol but not for R-propranolol binds similarly to a co-crystallized carazolol from the original structure (**Fig. 7B**). The top 1 pose of l-sotalol and top 2 pose of d-sotalol, with similar scores, have aligned their sulfonamide groups similarly (**Fig. 7B**). The top 1 pose for d-sotalol (not shown), scored considerable higher than the top 2 pose (−14.26 REU) and had flipped such that its propran-2-ylamino group was located where the methanesulfonamide position was for the top 2 pose.

**Figure 7.**
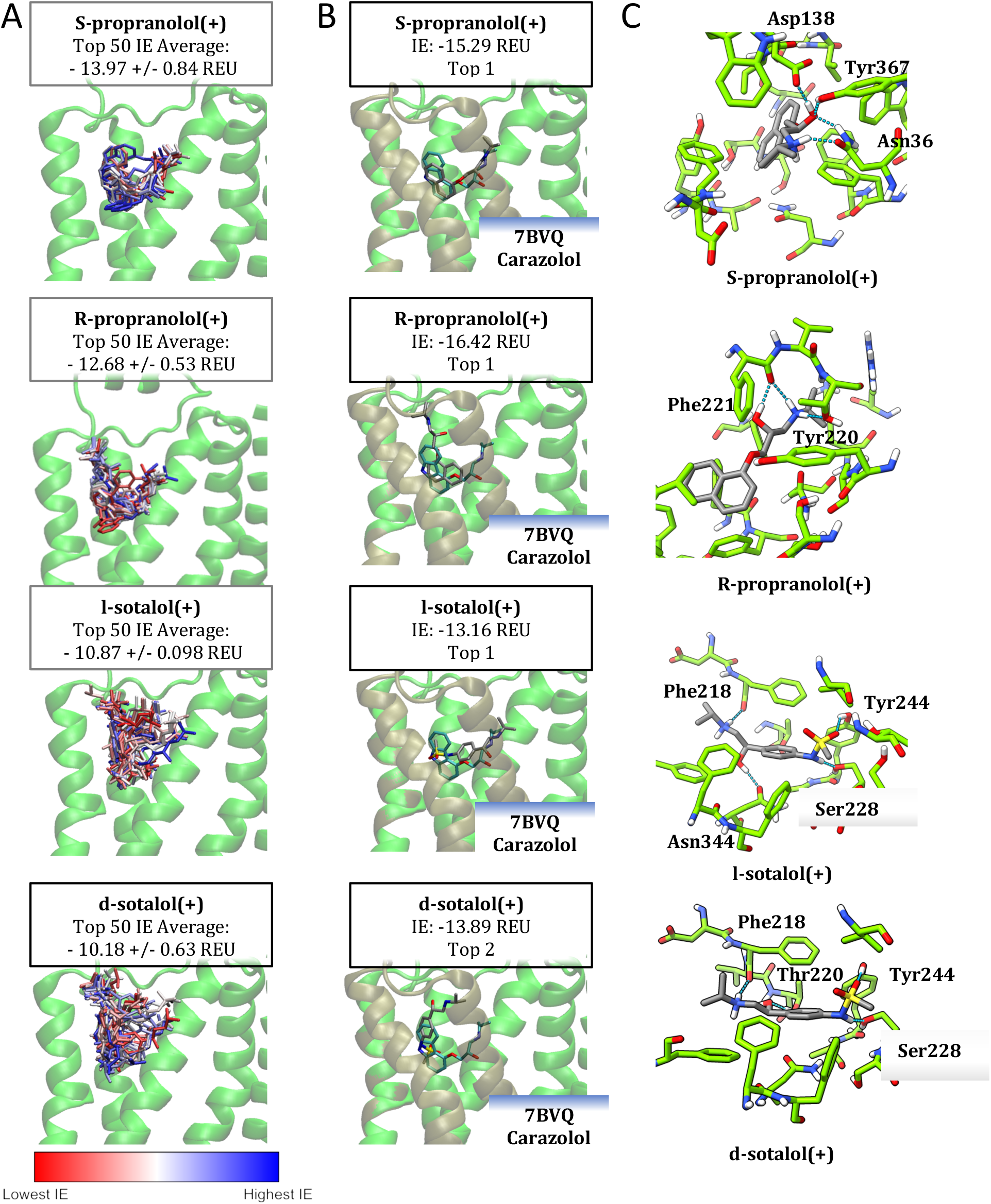
RosettaLigand docking stereoisomers of sotalol and propranolol into inactive-state human beta-1 adrenergic receptor. (A) Top fifty poses of cationic drug docking into the inactive human β_1_AR model colored from lowest interaction energy, IE (red, most favorable) to highest IE (blue, least favorable. (**B**) Candidate pose selected from top ten scoring poses compared to crystalline carazolol, a beta-blocker (cyan) as bound to the original template structural model (tan) PDB ID: 7BVQ. (**C**) PLIP contact analysis of the candidate pose for each drug with interacting residues labeled and H-bonds depicted by dotted blue lines.

We no longer observe participation of serine in the top poses of RS-propranolol docked to the inactive state; instead Tyr367 or Tyr220 participate in the binding (**Fig. 7C**). For sotalol, both candidate poses interact with Ser228, Thr220, and Phe218, with the distinction being the coordination of either Thr220 or Asn344 with the hydroxyl moiety of sotalol, a consequence of their chiral center inversion (**Fig. 7C**).

When overlaid, the ensembles of top fifty scoring poses for each drug diverge significantly in general shape and orientation for active β_1_AR model than those obtained from docking to the inactive state receptor model (**Fig. 6A & Fig. 7A**), suggesting that the volume available for the ligands is quite different. This may account for the different binding modalities observed in the sotalol ensembles. When docking to the inactive state, the sotalol adopt a linear orientation with the sulfonamide “digging” into the binding pocket (**Fig. 7A**). However, in the inactive state, there is a bifurcation in orientations that differ from the active-state modality (**Fig. 6A**).

We observe some distinctions in ensemble orientations when comparing the top fifty poses of beta-blocker docking against either turkey-derived β_1_AR homology model or the human-derived β_1_AR model. For instance R- and S-propranolol binding pose ensembles were tighter when bound to the human-derived model in comparison to the homology model, where the set of poses adopted a flatter, more distributed shape **(cf. Figs. 3A & 7A**).The top interfacial energy scores for either stereoisomer improved by about one Rosetta energy unit form their corresponding top poses against the β_1_AR homology model (cf.**Figure 3B** and **7B**). The top fifty poses of active-state propranolol binding β_1_AR are dominated by two observable orientations, with our docked R-propranolol seemingly favoring the binding orientation seen in the crystallographic pose of S-propranolol PDB ID 6PS5, and docked S-propranolol favoring the opposite orientation (**Fig. 7A**). The inactive-state β_1_AR - propranolol docking abolishes this pattern in the top 50 poses (**Fig. 7A**), and S-propranolol poses form a tight cloud around the orthosteric ligand binding pocket in contrast to R-propranolol, which exhibits a binding modality more like sotalol, interacting with the β_1_AR extracellular vestibule.

We also analyzed interacting β_1_AR residues for each top ligand binding pose as shown in panels C of **Figures 6 and 7**. One consistent pattern between the top poses of docked sotalol is the participation of Ser228 for both inactive and active states with the exception of active β_1_AR - d-sotalol interaction (**Fig. 6C & Fig. 7C**). In addition, Asn344 participates in active, but not inactive state β_1_AR binding of sotalol (**Fig. 6C & Fig. 7C**) Validating whether this pattern is indicative of a stereospecific, state-dependent interaction necessitates a more thorough examination of the full data set than is presented here. Lastly, it is of note that the preferential interaction of R-propranolol is consistent across β_1_AR model systems discussed here. R-propranolol has a more favorable interfacial score than S-propranolol when docked to the inactive human β_1_AR homology model based on turkey receptor structure (**Fig. 3B**) as well as new inactive-state (**Fig. 7B**) and active-state (**Fig. 6B**) β_1_AR models developed using human receptor structures.

## 4. Discussion

In this work, we constructed the first homology models of contiguous human β_1_AR in the active state in complex with the stimulatory G-protein and in the inactive state before the publication of human structures. We similarly developed models of active and inactive β_2_AR and active β_2_AR-G_s_ complex. We docked norepinephrine into the β_2_AR models and found the top poses to be in good agreement with a β_2_AR structure co-crystallized with epinephrine. Developed norepinephrine-bound active β_2_AR and β_2_AR-G_s_ structural models were successfully used for our recent all-atom MD simulation study,^17^ were we observed tighter binding of norepinephrine to β_2_AR-G_s_ compared to β_2_AR in agreement with experimental data, several alternative binding sites of the norepinephrine in the receptor cavity, observed interplay between different conformations of G_s_ and its partial dissociation from the receptor. That study validated our Rosetta modeling protocol.

Docking beta blockers dl-sotalol and RS-propranolol to the inactive homology model of human β_1_AR indicated more favorable energetic interactions for propranolol than sotalol, but no preference for a stereoisomer was observed. Subsequently simulating the docked stereoisomers of sotalol in multi-microsecond molecular dynamics simulations indicated that both binding poses were stable but did not differ significantly in their binding as could be expected based on experimental information. ^13^

Upon the release of human β_1_AR crystal structures, we developed new models of this receptor with an alternative protocol using the latest Rosetta Membrane Energy Function. In this case, the active-state human β_1_AR model interactions with beta blockers recapitulated the findings with the homology models. However, the average interfacial scores (interaction energies) for the top fifty poses against the inactive state suggest a slight preference for the l-sotalol and S-propranolol, ax expected.

This work can be improved upon through devising enhanced sampling molecular dynamics experiments that sample the energetics along a reaction coordinate that captures the complete transit of the drug from solvent into the binding pocket for reasons that are discussed below. To assess drug – protein interaction during drug entrance or egress may identify the determinizing factors for stereospecific drug binding. Drug flooding MD simulations ^12, 61^ can be also used identify reaction coordinates for drug access. Once a reaction coordinate is determined, enhanced sampling molecular dynamics such as umbrella sampling^62^ along with Hamiltonian replica exchange ^63^ or well-tempered metadynamics ^64^ should be sufficient for determining the free energy profiles and subsequently the dissociation constants for either stereoisomer of either drug^12, 65^. Alternatively, Gaussian accelerated MD simulations,^66^ weighted ensemble method^67^ or their combination^68^ can be used. These simulations will be explored in our follow-up studies.

### 4.1. The outer vestibule of the receptor as a possible mechanism of stereoselectivity

One reason that may account for the small difference between stereoisomer Rosetta energies is the possibility that a different portion of the protein is conveying selectivity. Dror et al. have shown in MD simulations that ligands encounter a large energy barrier when transitioning from the vestibule, an intermediate region between the extracellular space and the binding pocket, into the binding pocket^69^. This intermediary binding site may be more selective than the binding pocket itself, so while d-sotalol and l-sotalol are theoretically very stable when occupying the pocket, d-sotalol may simply have a more difficult time passing the vestibule.. It is of note that there are very few differences in sequence identity between the β_1_AR and β_2_ AR subtypes within the binding pocket itself; they are effectively identical in sequence within the pocket, with only a Phe/Tyr substitution^70^. Otherwise, the most proximal changes in the amino acid sequence reside at the edge of the pocket, near the vestibule^60^. Similar findings were observed to account for differences for NE and epinephrine binding to the β_1_AR and β_2_ AR in the recent study. ^34^

### 4.2. Regarding the selection of homology models and clustered models

Further iterations of relaxation of the top decoy of each candidate model may provide better criteria for decoy selection and provide a quantitative justification for selecting one cluster over another, as opposed to assuming that membrane-penetration by a protein chain is not feasible. Alternatively, we should consider using the most-likely decoy as opposed to the lowest energy decoy. This would mean considering the distribution of scores within a cluster and thus select multiple mean-value decoys from which a second round of clustering may be used assess the validity of each cluster in this new set of mean decoys. However, molecular dynamics samples conformations extensively. **Fig. S6** demonstrates how rapidly the ICL3 can rapidly adopt alternative conformations that would significantly increase protein root-mean-square deviation (RMSD) values, but at the same time it may still retain some secondary structure elements.

### 4.3. The possibility of allostery not captured by docking

It is important to acknowledge that the human β_1_AR homology model was derived from an isoprenaline-bound template that resembles an inactive structure. In other words, we docked antagonists into a formerly agonist-bound pocket. This may explain why propranolol failed to recapitulate the pose adopted in a known crystallographic binding pose of the ligand during docking (**Fig. 3**). This may indicate that suggests that perturbing local sidechain and backbone orientation is inadequate to capture beta-blocker-like interactions with the receptor. Larger backbone movements may be associated with facilitating inactivation within the binding pocket, movements that cannot be sampled by RosettaLigand with the present protocol. In the case of sotalol, it may be that the structure is flexible, but we also may have not captured the experimental binding pose in our ensembles. Future molecular dynamics simulations ought to examine receptor backbone reorganizations and potentially correlate such movements with drug orientation in the binding pocket and could be assessed using e.g. position and polar angle of the ligand binding orientation as we did in a recent study. ^12^

### 4.4. Alternative measures, a new hypothesis, and alternative sites of stereoselective ligand discrimination

One advantage of assessing stereoisomers is that one may appropriately compare the scores between them when evaluating drug docking. A more thorough examination of docking energetics, hydrogen bonding networks, and contact maps should be performed on the top 1 to 5 percent of binding poses to yield a more substantial dataset for such analysis: for instance, Smith and Meiler found in screening tests that within the top 1% of top-scoring poses, 21% of those poses will recapitulate a native binding pose when using the Rosetta Score function Talaris2014^71^, with this dropping off 6.9% for any of four assessed score functions^72^. This drops to 4.4% and 3.1% when using the top 5% or 10% of poses respectively^73^. This means when examining the top 5,000 poses of 50,000, we may conservatively estimate that 155 models will recapitulate native binding. If we select to 1%, we can expect 34 of 500 poses to be accurate representations of native binding; using Talaris2014 could raise this to one in five models. At that level of fidelity, introducing alternative quantities such as binding density, the ratio of binding energy to buried surface area, can provide new measures to support alternative candidate docked ligand conformations, or perhaps ligand clustering would be a sufficient to determine a candidate. Ultimately, determining the energetics and kinetics of the interactions based upon an identified reaction coordinate would circumvent the present inadequacies. Determining the dissociation constants of dl-sotalol against β_1_AR across multiple states using enhanced sampling methodologies discussed above can address the inadequacies in the present study or provide alternative hypotheses that may reveal allosteric interactions necessary for β_1_AR to discriminate between stereoisomers. One key consideration is whether access via the previously discussed extracellular vestibule above the binding pocket is stereospecific. Dror et al observed a metastable binding position for the beta blocker alprenolol occurring at the extracellular vestibule above the receptor;^69^ they concluded that the primary barrier to binding is this metastable site outside of the binding pocket. While our top fifty poses are located in the regions adjacent to the inner vestibule in varying orientations it is unclear whether we can readily observe this phenomenon from the present analysis (**Fig. 7A&B**). We would hypothesize that the molecular mechanism for stereospecific selectivity occurs at this vestibule rather than the binding pocket, and is perhaps state-dependent, given our present conclusions. Therefore, MD simulations should be conducted to search not only the absolute free-energy minimum within the binding pocket, but also for free energy minima that serve as a metastable state preceding interaction with the orthosteric binding pocket. Should atomistic molecular dynamics be unfeasible for assessing the along this access pathway, then adjusting the RosettaLigand docking window to target the extracellular vestibule warrants consideration. Capping this vestibule is the helix on the extracellular loop 2 (ECL2). To test this hypothesis, examination of ECL2 motility from previous simulations and assessing sequence conservation between receptors in relation to stereospecific ligand affinities would be a first step towards assessing extracellular portions of the receptor governing stereoselectivity of ligand binding.

## Acknowledgments

We would like to thank all members of the I.V., C.E.C. and V.Y.-Y. laboratories for helpful discussions. This work was supported by National Institutes of Health Common Fund Grant OT2OD026580 (to C.E.C. and I.V.), National Heart, Lung, and Blood Institute (NHLBI) grants R01HL128537, R01HL152681, and U01HL126273 (to C.E.C., V.Y.-Y. and I.V.), American Heart Association Career Development Award grant 19CDA34770101 (to I.V.), National Science Foundation travel grant 2032486 (to I.V.), UC Davis Department of Physiology and Membrane Biology Research Partnership Fund (to C.E.C. and I.V.) as well as UC Davis T32 Predoctoral Training in in Pharmacological Sciences fellowship supported in part by NHLBI Institutional Training Grant T32GM099608 (to J.R.D.D.). Computer allocations were provided through Extreme Science and Engineering Discovery Environment (XSEDE) grant MCB170095 (to I.V., C.E.C., and V.Y.-Y.), Texas Advanced Computing Center (TACC) Leadership Resource and Pathways Allocations MCB20010 (I.V., C.E.C., and V.Y.-Y.), Oracle for Research fellowship and cloud credits award (to I.V., C.E.C.), Pittsburgh Supercomputing Center (PSC) Anton 2 allocations PSCA17085P, MCB160089P, PSCA18077P, PSCA17085P, PSCA16108P (to I.V., C.E.C., and V.Y.-Y.). Anton 2 computer time was provided by the Pittsburgh Supercomputing Center (PSC) through Grant R01GM116961 from the National Institutes of Health. The Anton 2 machine at PSC was generously made available by D.E. Shaw Research.

## Supporting Information

Supporting information for this paper includes 1 Table (S1) and 6 Figures (S1-S6) providing additional details on beta-adrenergic receptor and G protein amino acid residue sequences, Rosetta structural modeling, RosettaLigand docking and molecular dynamics simulation results.

## References.

(1) Cannon, W. B. Bodily Changes in Pain, Hunger, Fear, And Rage: An Account of Recent Searches into the Function of Emotional Excitement; D. Appleton and Company, 1922.

(2) McCorry, L. K. Physiology of the autonomic nervous system. Am J Pharm Educ 2007, 71 (4), 78. DOI: 10.5688/aj710478 From NLM.

(3) Alshak, M. N.; J, M. D. Neuroanatomy, Sympathetic Nervous System. In StatPearls, StatPearls Publishing Copyright © 2022, StatPearls Publishing LLC., 2021.

(4) Ripplinger, C. M.; Noujaim, S. F.; Linz, D. The nervous heart. Prog Biophys Mol Biol 2016, 120 (1-3), 199–209. DOI: 10.1016/j.pbiomolbio.2015.12.015 From NLM. Goldberger, J. J.; Arora, R.; Buckley, U.; Shivkumar, K. Autonomic Nervous System Dysfunction: JACC Focus Seminar. *J Am Coll Cardiol* **2019**, *73* (10), 1189-1206. DOI: 10.1016/j.jacc.2018.12.064 PubMed.

(5) Grandi, E.; Ripplinger, C. M. Antiarrhythmic mechanisms of beta blocker therapy. Pharmacol Res 2019, 146, 104274–104274. DOI: 10.1016/j.phrs.2019.104274 PubMed.

(6) Lymperopoulos, A.; Rengo, G.; Koch, W. J. Adrenergic nervous system in heart failure: pathophysiology and therapy. Circ Res 2013, 113 (6), 739–753. DOI: 10.1161/circresaha.113.300308 From NLM. Bylund, D. B.; Eikenberg, D. C.; Hieble, J. P.; Langer, S. Z.; Lefkowitz, R. J.; Minneman, K. P.; Molinoff, P. B.; Ruffolo, R. R., Jr.; Trendelenburg, U. International Union of Pharmacology nomenclature of adrenoceptors. *Pharmacol Rev* **1994**, *46* (2), 121-136. From NLM.

(7) Woo, A. Y. H.; Xiao, R.-p. β-Adrenergic receptor subtype signaling in heart: from bench to bedside. Acta pharmacologica Sinica 2012, 33 (3), 335–341. DOI: 10.1038/aps.2011.201 PubMed.

(8) Wachter, S. B.; Gilbert, E. M. Beta-adrenergic receptors, from their discovery and characterization through their manipulation to beneficial clinical application. Cardiology 2012, 122 (2), 104–112. DOI: 10.1159/000339271 From NLM.

(9) Baker, J. G. The selectivity of beta-adrenoceptor antagonists at the human beta1, beta2 and beta3 adrenoceptors. Br J Pharmacol 2005, 144 (3), 317-322. DOI: 10.1038/sj.bjp.0706048 From NLM.

(10) Liu, X.; Luo, D.; Zhang, J.; Du, L. Distribution and relative expression of vasoactive receptors on arteries. Scientific Reports 2020, 10 (1), 15383. DOI: 10.1038/s41598-020-72352-5. Srinivasan, A. V. Propranolol: A 50-Year Historical Perspective. *Ann Indian Acad Neurol* **2019**, *22* (1), 21-26. DOI: 10.4103/aian.AIAN_201_18 PubMed.

(11) do Vale, G. T.; Ceron, C. S.; Gonzaga, N. A.; Simplicio, J. A.; Padovan, J. C. Three Generations of β-blockers: History, Class Differences and Clinical Applicability. Curr Hypertens Rev 2019, 15 (1), 22–31. DOI: 10.2174/1573402114666180918102735 From NLM.

(12) DeMarco, K. R.; Yang, P.-C.; Singh, V.; Furutani, K.; Dawson, J. R. D.; Jeng, M.-T.; Fettinger, J.; Bekker, S.; Ngo, V. A.; Noskov, S. Y.;, et al. Molecular determinants of pro-arrhythmia proclivity of d- and l-sotalol via a multi-scale modeling pipeline. Journal of Molecular and Cellular Cardiology 2021. DOI: 10.1016/j.yjmcc.2021.05.015.

(13) Funck-Brentano, C. Pharmacokinetic and pharmacodynamic profiles of d-sotalol and d, l-sotalol. European heart journal 1993, 14 (suppl_H), 30-35.

(14) Pratt, C. M.; Camm, A. J.; Cooper, W.; Friedman, P. L.; MacNeil, D. J.; Moulton, K. M.; Pitt, B.; Schwartz, P. J.; Veltri, E. P.; Waldo, A. L. Mortality in the Survival With ORal D-sotalol (SWORD) trial: why did patients die? Am J Cardiol 1998, 81 (7), 869–876. DOI: 10.1016/s0002-9149(98)00006-x From NLM.

(15) Mehvar, R.; Brocks, D. R. Stereospecific pharmacokinetics and pharmacodynamics of beta-adrenergic blockers in humans. J Pharm Pharm Sci 2001, 4 (2), 185–200. From NLM.

(16) Yao, X.; McIntyre, M. S.; Lang, D. G.; Song, I. H.; Becherer, J. D.; Hashim, M. A. Propranolol inhibits the human ether-a-go-go-related gene potassium channels. Eur J Pharmacol 2005, 519 (3), 208–211. DOI: 10.1016/j.ejphar.2005.05.010 From NLM. Dupuis, D. S.; Klaerke, D. A.; Olesen, S. P. Effect of beta-adrenoceptor blockers on human ether-a-go-go-related gene (HERG) potassium channels. *Basic Clin Pharmacol Toxicol* **2005**, *96* (2), 123-130. DOI: 10.1111/j.1742-7843.2005.pto960206.x From NLM. Chatzidou, S.; Kontogiannis, C.; Tsilimigras, D. I.; Georgiopoulos, G.; Kosmopoulos, M.; Papadopoulou, E.; Vasilopoulos, G.; Rokas, S. Propranolol Versus Metoprolol for Treatment of Electrical Storm in Patients With Implantable Cardioverter-Defibrillator. *J Am Coll Cardiol* **2018**, *71* (17), 1897-1906. DOI: 10.1016/j.jacc.2018.02.056 From NLM.

(17) Han, Y.; Dawson, J. R.; DeMarco, K. R.; Rouen, K. C.; Bekker, S.; Yarov-Yarovoy, V.; Clancy, C. E.; Xiang, Y. K.; Vorobyov, I. Elucidation of a dynamic interplay between a beta-2 adrenergic receptor, its agonist, and stimulatory G protein. Proceedings of the National Academy of Sciences 2023, 120 (10), e2215916120.

(18) Ring, A. M.; Manglik, A.; Kruse, A. C.; Enos, M. D.; Weis, W. I.; Garcia, K. C.; Kobilka, B. K. Adrenaline-activated structure of β2-adrenoceptor stabilized by an engineered nanobody. Nature 2013, 502 (7472), 575–579. DOI: 10.1038/nature12572.

(19) Rasmussen, S. G.; Choi, H. J.; Rosenbaum, D. M.; Kobilka, T. S.; Thian, F. S.; Edwards, P. C.; Burghammer, M.; Ratnala, V. R.; Sanishvili, R.; Fischetti, R. F.;, et al. Crystal structure of the human beta2 adrenergic G-protein-coupled receptor. Nature 2007, 450 (7168), 383–387. DOI: 10.1038/nature06325 From NLM.

(20) Lomize, M. A.; Pogozheva, I. D.; Joo, H.; Mosberg, H. I.; Lomize, A. L. OPM database and PPM web server: resources for positioning of proteins in membranes. Nucleic Acids Res 2012, 40 (Database issue), D370-D376. DOI: 10.1093/nar/gkr703 PubMed.

(21) Pettersen, E. F.; Goddard, T. D.; Huang, C. C.; Couch, G. S.; Greenblatt, D. M.; Meng, E. C.; Ferrin, T. E. UCSF Chimera--a visualization system for exploratory research and analysis. J Comput Chem 2004, 25 (13), 1605–1612. DOI: 10.1002/jcc.20084 From NLM.

(22) Wang, C.; Bradley, P.; Baker, D. Protein-protein docking with backbone flexibility. J Mol Biol 2007, 373 (2), 503–519. DOI: 10.1016/j.jmb.2007.07.050 From NLM. Canutescu, A. A.; Dunbrack, R. L., Jr. Cyclic coordinate descent: A robotics algorithm for protein loop closure. *Protein Sci* **2003**, *12* (5), 963-972. DOI: 10.1110/ps.0242703 From NLM.

(23) Consortium, T. U. UniProt: the universal protein knowledgebase in 2021. Nucleic Acids Res 2020, 49 (D1), D480–D489. DOI: 10.1093/nar/gkaa1100 (acccessed 4/19/2022).

(24) Waterhouse, A. M.; Procter, J. B.; Martin, D. M. A.; Clamp, M.; Barton, G. J. Jalview Version 2—a multiple sequence alignment editor and analysis workbench. Bioinformatics 2009, 25 (9), 1189–1191. DOI: 10.1093/bioinformatics/btp033 (acccessed 5/11/2022).

(25) Alford, R. F.; Fleming, P. J.; Fleming, K. G.; Gray, J. J. Protein Structure Prediction and Design in a Biologically Realistic Implicit Membrane. Biophys J 2020, 118 (8), 2042–2055. DOI: 10.1016/j.bpj.2020.03.006 From NLM.

(26) Song, Y.; DiMaio, F.; Wang, R. Y.; Kim, D.; Miles, C.; Brunette, T.; Thompson, J.; Baker, D. High-resolution comparative modeling with RosettaCM. Structure 2013, 21 (10), 1735–1742. DOI: 10.1016/j.str.2013.08.005. Alford, R. F.; Leaver-Fay, A.; Jeliazkov, J. R.; O’Meara, M. J.; DiMaio, F. P.; Park, H.; Shapovalov, M. V.; Renfrew, P. D.; Mulligan, V. K.; Kappel, K.; et al. The Rosetta All-Atom Energy Function for Macromolecular Modeling and Design. *J Chem Theory Comput* **2017**, *13* (6), 3031-3048. DOI: 10.1021/acs.jctc.7b00125 From NLM.

(27) Yarov-Yarovoy, V.; Schonbrun, J.; Baker, D. Multipass membrane protein structure prediction using Rosetta. Proteins 2006, 62 (4), 1010–1025. DOI: 10.1002/prot.20817 From NLM.

(28) Bonneau, R.; Strauss, C. E. M.; Rohl, C. A.; Chivian, D.; Bradley, P.; Malmström, L.; Robertson, T.; Baker, D. De Novo Prediction of Three-dimensional Structures for Major Protein Families. Journal of Molecular Biology 2002, 322 (1), 65–78. DOI: 10.1016/S0022-2836(02)00698-8.

(29) Conway, P.; Tyka, M. D.; DiMaio, F.; Konerding, D. E.; Baker, D. Relaxation of backbone bond geometry improves protein energy landscape modeling. Protein Sci 2014, 23 (1), 47–55. DOI: 10.1002/pro.2389 From NLM.

(30) Warne, T.; Edwards, P. C.; Doré, A. S.; Leslie, A. G. W.; Tate, C. G. Molecular basis for high-affinity agonist binding in GPCRs. Science 2019, 364 (6442), 775–778. DOI: 10.1126/science.aau5595 From NLM.

(31) Sievers, F.; Wilm, A.; Dineen, D.; Gibson, T. J.; Karplus, K.; Li, W.; Lopez, R.; McWilliam, H.; Remmert, M.; Söding, J.;, et al. Fast, scalable generation of high-quality protein multiple sequence alignments using Clustal Omega. Molecular Systems Biology 2011, 7 (1), 539. DOI: 10.1038/msb.2011.75. Madeira, F.; Pearce, M.; Tivey, A. R. N.; Basutkar, P.; Lee, J.; Edbali, O.; Madhusoodanan, N.; Kolesnikov, A.; Lopez, R. Search and sequence analysis tools services from EMBL-EBI in 2022. *Nucleic Acids Res* **2022**. DOI: 10.1093/nar/gkac240 (acccessed 5/10/2022).

(32) Alford, R. F.; Fleming, P. J.; Fleming, K. G.; Gray, J. J. Protein Structure Prediction and Design in a Biologically Realistic Implicit Membrane. Biophysical Journal 2020, 118 (8), 2042–2055. DOI: 10.1016/j.bpj.2020.03.006.

(33) Warne, T.; Moukhametzianov, R.; Baker, J. G.; Nehmé, R.; Edwards, P. C.; Leslie, A. G.; Schertler, G. F.; Tate, C. G. The structural basis for agonist and partial agonist action on a β(1)-adrenergic receptor. Nature 2011, 469 (7329), 241–244. DOI: 10.1038/nature09746 From NLM.

(34) Xu, X.; Kaindl, J.; Clark, M. J.; Hübner, H.; Hirata, K.; Sunahara, R. K.; Gmeiner, P.; Kobilka, B. K.; Liu, X. Binding pathway determines norepinephrine selectivity for the human β(1)AR over β(2)AR. Cell Res 2021, 31 (5), 569–579. DOI: 10.1038/s41422-020-00424-2 From NLM.

(35) Mandell, D. J.; Coutsias, E. A.; Kortemme, T. Sub-angstrom accuracy in protein loop reconstruction by robotics-inspired conformational sampling. Nature Methods 2009, 6 (8), 551–552. DOI: 10.1038/nmeth0809-551.

(36) Meiler, J.; Baker, D. ROSETTALIGAND: protein-small molecule docking with full side-chain flexibility. Proteins 2006, 65 (3), 538–548. DOI: 10.1002/prot.21086 From NLM. Davis, I. W.; Baker, D. RosettaLigand docking with full ligand and receptor flexibility. *J Mol Biol* **2009**, *385* (2), 381-392. DOI: 10.1016/j.jmb.2008.11.010 From NLM. Davis, I. W.; Raha, K.; Head, M. S.; Baker, D. Blind docking of pharmaceutically relevant compounds using RosettaLigand. *Protein Sci* **2009**, *18* (9), 1998-2002. DOI: 10.1002/pro.192 From NLM.

(37) Hawkins, P. C.; Skillman, A. G.; Warren, G. L.; Ellingson, B. A.; Stahl, M. T. Conformer generation with OMEGA: algorithm and validation using high quality structures from the Protein Databank and Cambridge Structural Database. J Chem Inf Model 2010, 50 (4), 572–584. DOI: 10.1021/ci100031x From NLM.

(38) Salentin, S.; Schreiber, S.; Haupt, V. J.; Adasme, M. F.; Schroeder, M. PLIP: fully automated protein–ligand interaction profiler. Nucleic Acids Res 2015, 43 (W1), W443–W447. DOI: 10.1093/nar/gkv315 (acccessed 5/10/2022).

(39) Jo, S.; Lim, J. B.; Klauda, J. B.; Im, W. CHARMM-GUI Membrane Builder for Mixed Bilayers and Its Application to Yeast Membranes. Biophysical Journal 2009, 97 (1), 50–58. DOI: 10.1016/j.bpj.2009.04.013. Lee, J.; Cheng, X.; Swails, J. M.; Yeom, M. S.; Eastman, P. K.; Lemkul, J. A.; Wei, S.; Buckner, J.; Jeong, J. C.; Qi, Y.; et al. CHARMM-GUI Input Generator for NAMD, GROMACS, AMBER, OpenMM, and CHARMM/OpenMM Simulations Using the CHARMM36 Additive Force Field. *Journal of Chemical Theory and Computation* **2016**, *12* (1), 405-413. DOI: 10.1021/acs.jctc.5b00935. Wu, E. L.; Cheng, X.; Jo, S.; Rui, H.; Song, K. C.; Dávila-Contreras, E. M.; Qi, Y.; Lee, J.; Monje-Galvan, V.; Venable, R. M.; et al. CHARMM-GUI Membrane Builder toward realistic biological membrane simulations. *J Comput Chem* **2014**, *35* (27), 1997-2004. DOI: 10.1002/jcc.23702 From NLM. Jo, S.; Kim, T.; Im, W. Automated builder and database of protein/membrane complexes for molecular dynamics simulations. *PLoS One* **2007**, *2* (9), e880-e880. DOI: 10.1371/journal.pone.0000880 PubMed. Lee, J.; Patel, D. S.; Ståhle, J.; Park, S. J.; Kern, N. R.; Kim, S.; Lee, J.; Cheng, X.; Valvano, M. A.; Holst, O.; et al. CHARMM-GUI Membrane Builder for Complex Biological Membrane Simulations with Glycolipids and Lipoglycans. *J Chem Theory Comput* **2019**, *15* (1), 775-786. DOI: 10.1021/acs.jctc.8b01066 From NLM.

(40) Brooks, B. R.; Brooks, C. L., 3rd; Mackerell, A. D., Jr.; Nilsson, L.; Petrella, R. J.; Roux, B.; Won, Y.; Archontis, G.; Bartels, C.; Boresch, S.; et al. CHARMM: the biomolecular simulation program. Journal of computational chemistry 2009, 30 (10), 1545–1614. DOI: 10.1002/jcc.21287 PubMed.

(41) Dror, R. O.; Mildorf, T. J.; Hilger, D.; Manglik, A.; Borhani, D. W.; Arlow, D. H.; Philippsen, A.; Villanueva, N.; Yang, Z.; Lerch, M. T.;, et al. Structural basis for nucleotide exchange in heterotrimeric G proteins. Science 2015, 348 (6241), 1361–1365. DOI: doi:10.1126/science.aaa5264.

(42) Huang, J.; Rauscher, S.; Nawrocki, G.; Ran, T.; Feig, M.; de Groot, B. L.; Grubmüller, H.; MacKerell Jr, A. D. CHARMM36m: an improved force field for folded and intrinsically disordered proteins. Nature methods 2017, 14 (1), 71.

(43) Klauda, J. B.; Venable, R. M.; Freites, J. A.; O’Connor, J. W.; Tobias, D. J.; Mondragon-Ramirez, C.; Vorobyov, I.; MacKerell, A. D.; Pastor, R. W. Update of the CHARMM All-Atom Additive Force Field for Lipids: Validation on Six Lipid Types. The Journal of Physical Chemistry B 2010, 114 (23), 7830–7843. DOI: 10.1021/jp101759q.

(44) Yoo, J.; Aksimentiev, A. New tricks for old dogs: improving the accuracy of biomolecular force fields by pair-specific corrections to non-bonded interactions. Physical Chemistry Chemical Physics 2018, 20 (13), 8432–8449.

(45) Jorgensen, W. L.; Chandrasekhar, J.; Madura, J. D.; Impey, R. W.; Klein, M. L. Comparison of simple potential functions for simulating liquid water. The Journal of Chemical Physics 1983, 79 (2), 926–935. DOI: 10.1063/1.445869.

(46) Vanommeslaeghe, K.; Hatcher, E.; Acharya, C.; Kundu, S.; Zhong, S.; Shim, J.; Darian, E.; Guvench, O.; Lopes, P.; Vorobyov, I.;, et al. CHARMM general force field: A force field for drug-like molecules compatible with the CHARMM all-atom additive biological force fields. J Comput Chem 2010, 31 (4), 671–690. DOI: 10.1002/jcc.21367 From NLM.

(47) Vanommeslaeghe, K.; MacKerell Jr, A. D. Automation of the CHARMM General Force Field (CGenFF) I: bond perception and atom typing. J. Chem. Inf. Model. 2012, 52, 3144–3154. Vanommeslaeghe, K.; Raman, E. P.; MacKerell, A. D. Automation of the CHARMM General Force Field (CGenFF) II: Assignment of Bonded Parameters and Partial Atomic Charges. *Journal of Chemical Information and Modeling* **2012**, *52* (12), 3155-3168. DOI: 10.1021/ci3003649.

(48) Mayne, C. G.; Saam, J.; Schulten, K.; Tajkhorshid, E.; Gumbart, J. C. Rapid parameterization of small molecules using the force field toolkit. Journal of computational chemistry 2013, 34 (32), 2757–2770.

(49) Humphrey, W.; Dalke, A.; Schulten, K. VMD: visual molecular dynamics. J. Mol. Graphics 1996, 14 (1), 33.

(50) Frisch, M. J.; Trucks, G. W.; Schlegel, H. B.; Scuseria, G. E.; Robb, M. A.; Cheeseman, J. R.; Zakrzewski, V. G.; Montgomery, J. A.; Stratmann, R. E.; Burant, J. C.;, et al. *Gaussian 98*; 1998. Mayne, C. G.; Saam, J.; Schulten, K.; Tajkhorshid, E.; Gumbart, J. C. Rapid parameterization of small molecules using the Force Field Toolkit. J Comput Chem 2013, 34 (32), 2757–2770. DOI: 10.1002/jcc.23422 From NLM.

(51) DeMarco, K. R.; Bekker, S.; Clancy, C. E.; Noskov, S. Y.; Vorobyov, I. Digging into Lipid Membrane Permeation for Cardiac Ion Channel Blocker d-Sotalol with All-Atom Simulations. Frontiers in pharmacology 2018, 9, 26–26. DOI: 10.3389/fphar.2018.00026 PubMed.

(52) Phillips, J. C.; Braun, R.; Wang, W.; Gumbart, J.; Tajkhorshid, E.; Villa, E.; Chipot, C.; Skeel, R. D.; Kalé, L.; Schulten, K. Scalable molecular dynamics with NAMD. J Comput Chem 2005, 26 (16), 1781–1802. DOI: 10.1002/jcc.20289 From NLM.

(53) Nosé, S. A unified formulation of the constant temperature molecular dynamics methods. The Journal of Chemical Physics 1984, 81 (1), 511–519. DOI: 10.1063/1.447334. Hoover, W. G. Canonical dynamics: Equilibrium phase-space distributions. *Physical Review A* **1985**, *31* (3), 1695-1697.

(54) Feller, S. E.; Zhang, Y.; Pastor, R. W.; Brooks, B. R. Constant pressure molecular dynamics simulation: The Langevin piston method. The Journal of Chemical Physics 1995, 103 (11), 4613–4621. DOI: 10.1063/1.470648.

(55) Darden, T.; York, D.; Pedersen, L. Particle mesh Ewald: An N⋅log(N) method for Ewald sums in large systems. The Journal of Chemical Physics 1993, 98 (12), 10089–10092. DOI: 10.1063/1.464397.

(56) Ryckaert, J.-P.; Ciccotti, G.; Berendsen, H. J. C. Numerical integration of the cartesian equations of motion of a system with constraints: molecular dynamics of n-alkanes. Journal of Computational Physics 1977, 23 (3), 327–341. DOI: 10.1016/0021-9991(77)90098-5.

(57) Shaw, D. E.; Dror, R. O.; Salmon, J. K.; Grossman, J. P.; Mackenzie, K. M.; Bank, J. A.; Young, C.; Deneroff, M. M.; Batson, B.; Bowers, K. J.;, et al. Millisecond-scale molecular dynamics simulations on Anton. In Proceedings of the Conference on High Performance Computing Networking, Storage and Analysis, Portland, Oregon; 2009. Shaw, D. E.; Maragakis, P.; Lindorff-Larsen, K.; Piana, S.; Dror, R. O.; Eastwood, M. P.; Bank, J. A.; Jumper, J. M.; Salmon, J. K.; Shan, Y.; et al. Atomic-Level Characterization of the Structural Dynamics of Proteins. *Science* **2010**, *330* (6002), 341-346. DOI: doi:10.1126/science.1187409.

(58) Ishchenko, A.; Stauch, B.; Han, G. W.; Batyuk, A.; Shiriaeva, A.; Li, C.; Zatsepin, N.; Weierstall, U.; Liu, W.; Nango, E.;, et al. Toward G protein-coupled receptor structure-based drug design using X-ray lasers. IUCrJ 2019, 6 (Pt 6), 1106–1119. DOI: 10.1107/s2052252519013137 From NLM.

(59) Ballesteros, J. A.; Weinstein, H. Integrated methods for the construction of three-dimensional models and computational probing of structure-function relations in G protein-coupled receptors. In Methods in Neurosciences, Sealfon, S. C. Ed.; Vol. 25; Academic Press, 1995; pp 366–428.

(60) Wu, Y.; Zeng, L.; Zhao, S. Ligands of Adrenergic Receptors: A Structural Point of View. Biomolecules 2021, 11 (7). DOI: 10.3390/biom11070936.

(61) Boiteux, C.; Vorobyov, I.; French, R. J.; French, C.; Yarov-Yarovoy, V.; Allen, T. W. Local anesthetic and antiepileptic drug access and binding to a bacterial voltage-gated sodium channel. Proc Natl Acad Sci U S A 2014, 111 (36), 13057–13062. DOI: 10.1073/pnas.1408710111.

(62) Torrie, G. M.; Valleau, J. P. Nonphysical sampling distributions in Monte Carlo free-energy estimation: Umbrella sampling. Journal of Computational Physics 1977, 23 (2), 187–199. DOI: 10.1016/0021-9991(77)90121-8.

(63) Jiang, W.; Luo, Y.; Maragliano, L.; Roux, B. Calculation of Free Energy Landscape in Multi-Dimensions with Hamiltonian-Exchange Umbrella Sampling on Petascale Supercomputer. Journal of Chemical Theory and Computation 2012, 8 (11), 4672–4680. DOI: 10.1021/ct300468g.

(64) Barducci, A.; Bussi, G.; Parrinello, M. Well-tempered metadynamics: a smoothly converging and tunable free-energy method. Physical review letters 2008, 100 (2), 020603.

(65) Yang, P. C.; DeMarco, K. R.; Aghasafari, P.; Jeng, M. T.; Dawson, J. R.; Bekker, S.; Noskov, S.; Yarov-Yarovoy, V.; Vorobyov, I.; Clancy, C. E. A Computational Pipeline to Predict Cardiotoxicity:From the Atom to the Rhythm. Circ Res 2020, 126, 947–964. DOI: 10.1161/CIRCRESAHA.119.316404.

(66) Miao, Y.; Bhattarai, A.; Wang, J. Ligand Gaussian accelerated molecular dynamics (LiGaMD): Characterization of ligand binding thermodynamics and kinetics. Journal of chemical theory and computation 2020, 16 (9), 5526–5547.

(67) Copperman, J.; Zuckerman, D. M. Accelerated Estimation of Long-Timescale Kinetics from Weighted Ensemble Simulation via Non-Markovian “Microbin” Analysis. J Chem Theory Comput 2020, 16 (11), 6763–6775. DOI: 10.1021/acs.jctc.0c00273.

(68) Ahn, S. H.; Ojha, A. A.; Amaro, R. E.; McCammon, J. A. Gaussian-Accelerated Molecular Dynamics with the Weighted Ensemble Method: A Hybrid Method Improves Thermodynamic and Kinetic Sampling. J Chem Theory Comput 2021, 17 (12), 7938–7951. DOI: 10.1021/acs.jctc.1c00770.

(69) Dror, R. O.; Pan, A. C.; Arlow, D. H.; Borhani, D. W.; Maragakis, P.; Shan, Y.; Xu, H.; Shaw, D. E. Pathway and mechanism of drug binding to G-protein-coupled receptors. Proceedings of the National Academy of Sciences 2011, 108 (32), 13118–13123. DOI: doi:10.1073/pnas.1104614108.

(70) Hedderich, J. B.; Persechino, M.; Becker, K.; Heydenreich, F. M.; Gutermuth, T.; Bouvier, M.; Bünemann, M.; Kolb, P. The pocketome of G-protein-coupled receptors reveals previously untargeted allosteric sites. Nature Communications 2022, 13 (1), 2567. DOI: 10.1038/s41467-022-29609-6.

(71) Leaver-Fay, A.; O’Meara, M. J.; Tyka, M.; Jacak, R.; Song, Y.; Kellogg, E. H.; Thompson, J.; Davis, I. W.; Pache, R. A.; Lyskov, S.;, et al. Scientific benchmarks for guiding macromolecular energy function improvement. Methods in enzymology 2013, 523, 109–143. DOI: 10.1016/B978-0-12-394292-0.00006-0 PubMed. O’Meara, M. J.; Leaver-Fay, A.; Tyka, M. D.; Stein, A.; Houlihan, K.; DiMaio, F.; Bradley, P.; Kortemme, T.; Baker, D.; Snoeyink, J.; et al. Combined Covalent-Electrostatic Model of Hydrogen Bonding Improves Structure Prediction with Rosetta. *Journal of Chemical Theory and Computation* **2015**, *11* (2), 609-622. DOI: 10.1021/ct500864r.

(72) Smith, S. T.; Meiler, J. Assessing multiple score functions in Rosetta for drug discovery. PLoS One 2020, 15 (10), e0240450–e0240450. DOI: 10.1371/journal.pone.0240450 PubMed.

(73) Leman, J. K.; Weitzner, B. D.; Lewis, S. M.; Adolf-Bryfogle, J.; Alam, N.; Alford, R. F.; Aprahamian, M.; Baker, D.; Barlow, K. A.; Barth, P.;, et al. Macromolecular modeling and design in Rosetta: recent methods and frameworks. Nat Methods 2020, 17 (7), 665–680. DOI: 10.1038/s41592-020-0848-2.

