## Supplemental table and figures for "Modeling Stereospecific Drug Interactions with Beta-Adrenergic Receptors"

**Table S1. Restraint regime for extended molecular dynamics equilibration.**

| Stage | Steps | Duration(ns) | Restraint (kcal/mol/Å$^2$) |
|---|---|---|---|
| 1 | 7.1 - 7.4 | 4 | 1.0 for all backbone Cα carbons. |
| 2 | 7.5 - 7.10 | 6 | 1.0 for backbone Cα carbons not modeled *de novo*. |
| 3 | 7.11 - 7.15 | 5 | 0.5 for backbone Cα carbons not modeled *de novo*. |
| 4 | 7.16 – 7.20 | 5 | 0.25 for backbone Cα carbons not modeled *de novo*. |
| 5 | 7.21 – 7.30 | 10 | 0.1 for backbone Cα carbons not modeled *de novo*. |
| 6 | 7.31 – 7.40 | 10 | 0.5 for backbone Cα carbons within 3.5 Å of docked or crystallographic ligand. |
| 7 | 7.41 – 90 | 50 | No restraints. |

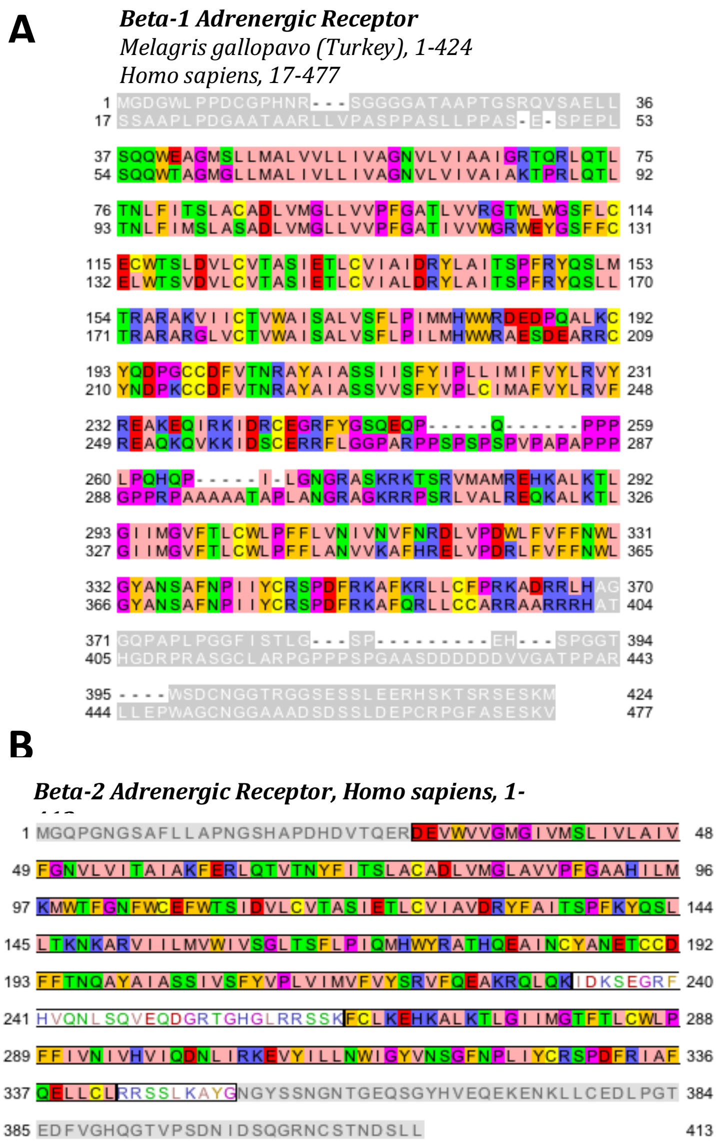


**<u>Figure S1.</u> Amino acid residue sequences of the Beta-1 Adrenergic Receptor ($\beta_1$AR) and Beta-2 Adrenergic Receptor ($\beta_2$AR).** (**A**) Pairwise sequence alignment of the amino acid residue sequences for $\beta_1$AR genes for turkey (above) and human (below) visualized using Jalview. Residues in ”Zappo” color scheme are modeled whereas grey-colored residues were omitted from modeling. The intracellular loop 3 (ICL3) for the human protein sequence spans residues 256 to 314 and was modelled *de novo*. *De novo* modeling of the C-terminus spans residues 342 to 350. (**B**) The $\beta_2$AR sequence included in the template model(”Zappo” coloring). Regions that were modeled *de novo* are denoted with colored text and white backgrounds: the ICL3 spanning residues 233 to 263 and the C-terminus spanning residues 342 to 351.

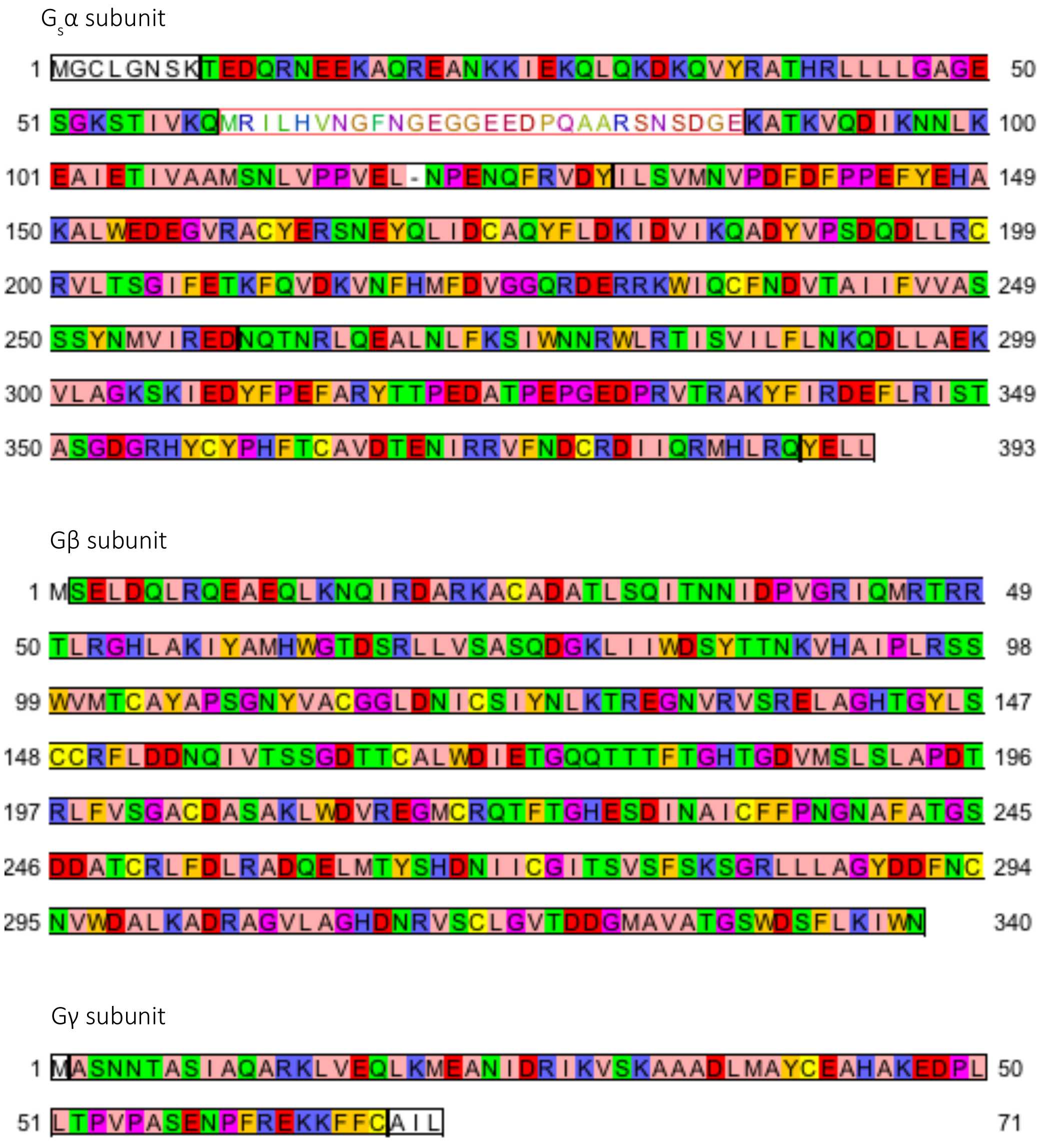


**<u>Figure S2.</u> Amino acid residue sequences of G protein subunits Gsα, Gβ, and Gγ.** All three sequences are imaged in "Zappo" coloration using Jalview. Sequences depicted in black and white including, N-terminal Met, which genetically encoded, were omitted from unresolved in the template model. *Gsα Subunit.* Residues 61 to 87 of Gsα were modeled *de novo* using RosettaCM and are depicted in colored text with white background. Residue 72 of Gsα ("-") is Ala. *Gβ Subunit.* All residues of the gene Gγ Subunit excluding N-terminal Methionine are modeled. Residues 62 to 68 of Gγ are modeled de novo and were not resolved in PDB ID 3SN6.

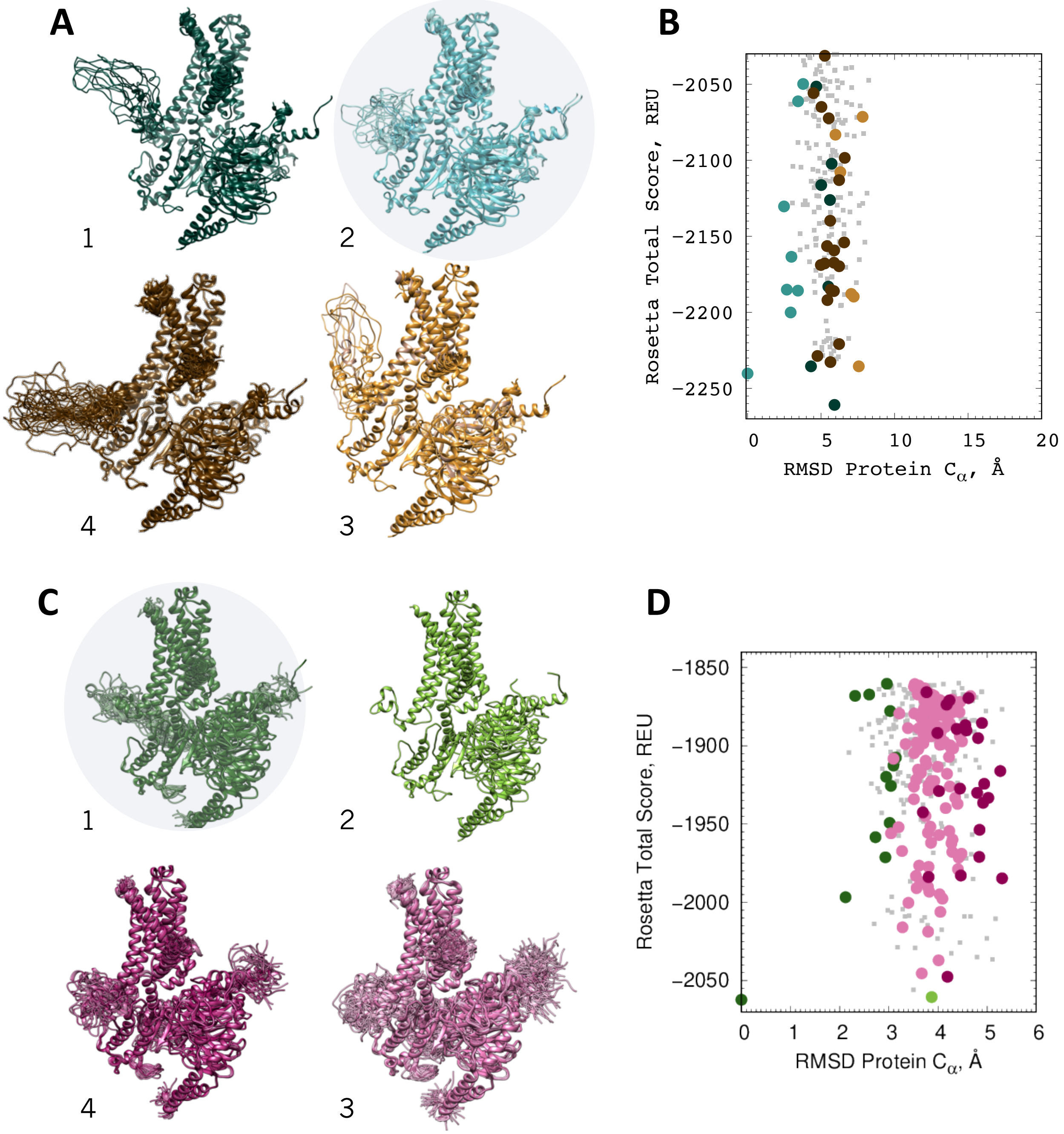


**<u>Figure S3</u>: Clustering analysis of active $\beta_1$AR/$G_s$ and $\beta_2$AR/$G_s$ models developed using RosettaCM protocol.** (**A**) Top four clusters for $\beta_1$AR/$G_s$ demonstrating that gross orientation of the $\beta_1$AR ICL3 (shown as a protrusion in the left central portion for each model) governs clustering. The candidate model is selected from cluster 2 (circled). (**B**) Graph of total Rosetta energy score for each clustered $\beta_1$AR decoy according to root-mean-square deviation (RMSD) from the candidate model. Top four clusters are colored according to their image on the left. (**C**) Top four clusters for $\beta_2$AR/$G_s$ demonstrating tighter convergence about the shorter ICL3 (small protrusion on the left for each model). The candidate model is selected from cluster 1 (circled). **D**) Graph of total Rosetta energy score for each clustered $\beta_2$AR decoy according to RMSD from the candidate model. Top four clusters are colored according to their images on the left.

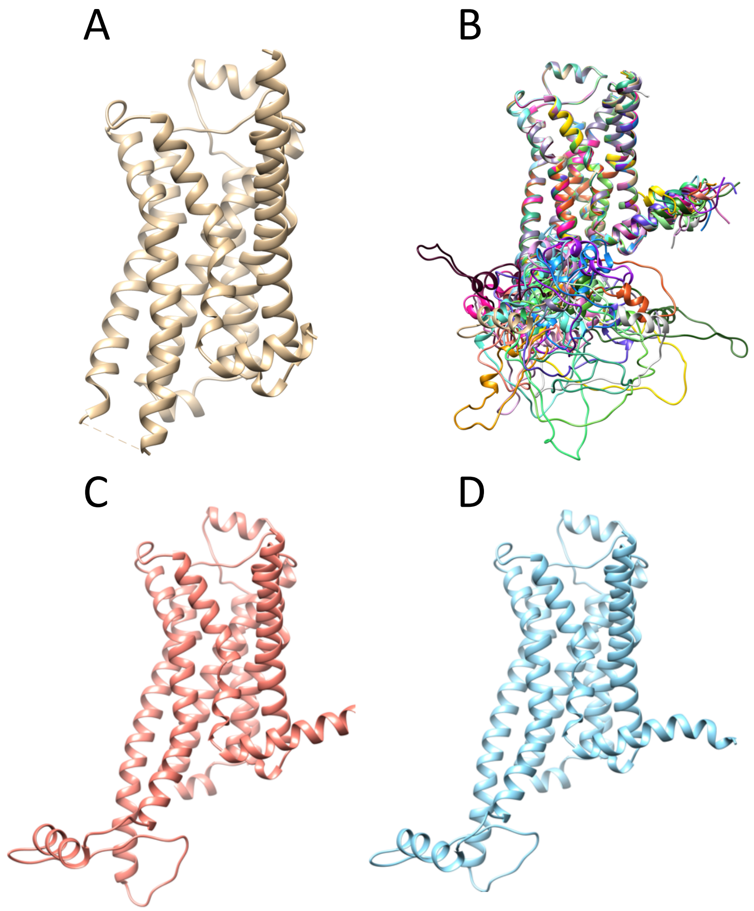


**<u>Figure S4.</u> Stages of homology modeling for inactive human $\beta_1$AR from turkey $\beta_1$AR (PDB ID: 2Y03).** (**A**) The template PDB after removal of ligands isoprenaline, cholesterol hemisuccinate, and HEGA-10**.** **(B**) Steric clashes prohibited incorporating a G protein into modeling; In the absence of the G-protein, the top decoys of the top 20 clusters demonstrate poor convergence when overlaid. (**C**) Candidate model selected from 10,000 decoy models. (**D**) Candidate model after restrained relaxation step.

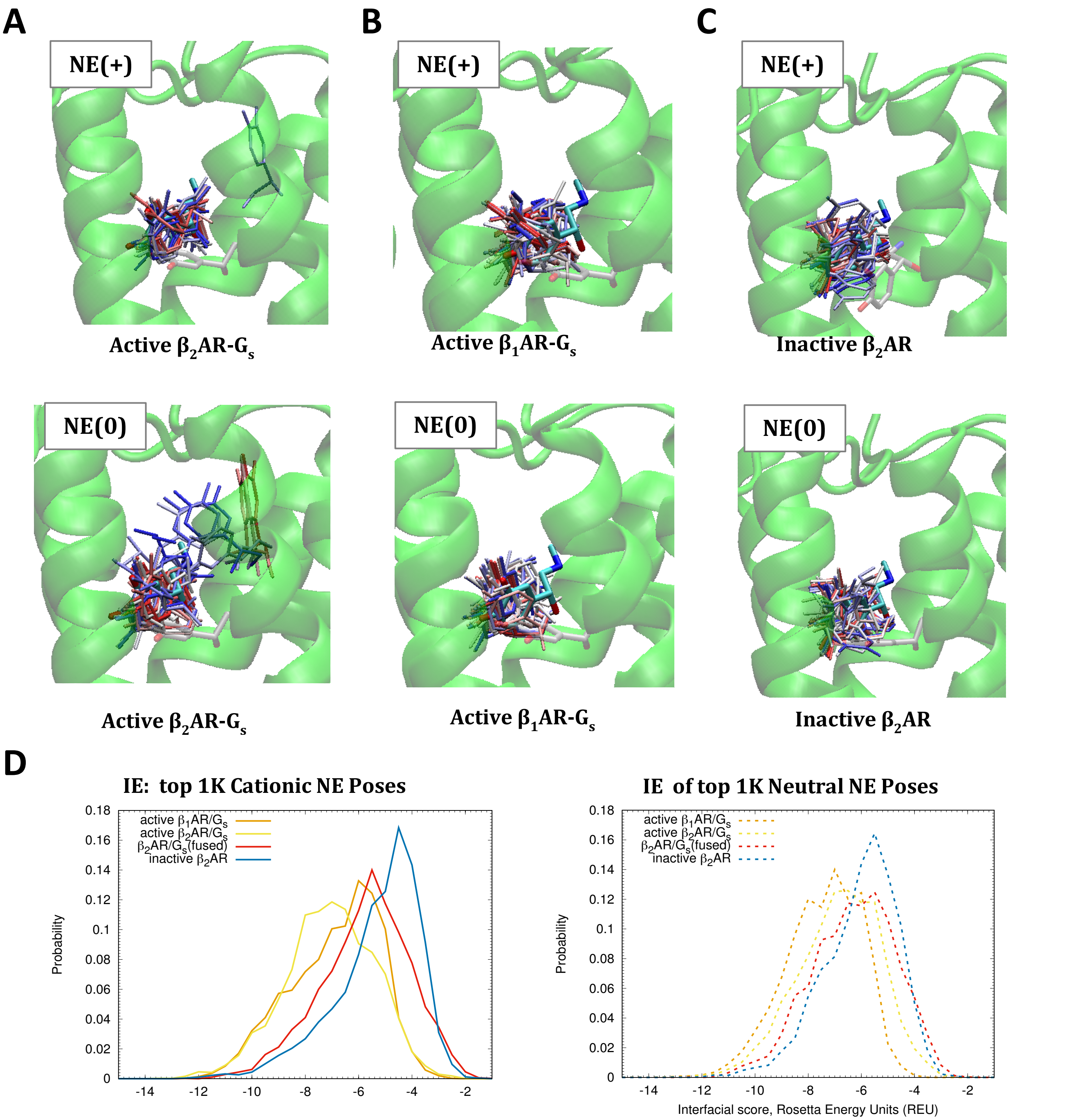


**<u>Figure S5.</u> RosettaLigand docking of norepinephrine into RosettaCM-derived $\beta_1$AR and $\beta_2$AR models .** (**A-C**) Top fifty poses of cationic (top) and neutral (bottom) norepinephrine (NE) into (**A**) active $\beta_2$AR-G$_s$, (**B**) active $\beta_1$AR-G$_s$, (**C**) inactive $\beta_2$AR homology models. Crystalized epinephrine of PDB 4LDO is depicted in cyan after alignment with the candidate models. (**D**) Probability distributions of Rosetta interfacial scores or interaction energies (IE)_for docked cationic (left) or neutral (right) norepinephrine against active $\beta_2$AR, inactive $\beta_2$AR, active $\beta_1$AR models as well as G$_s$-fused $\beta_2$AR structure (PDB ID 6E67).

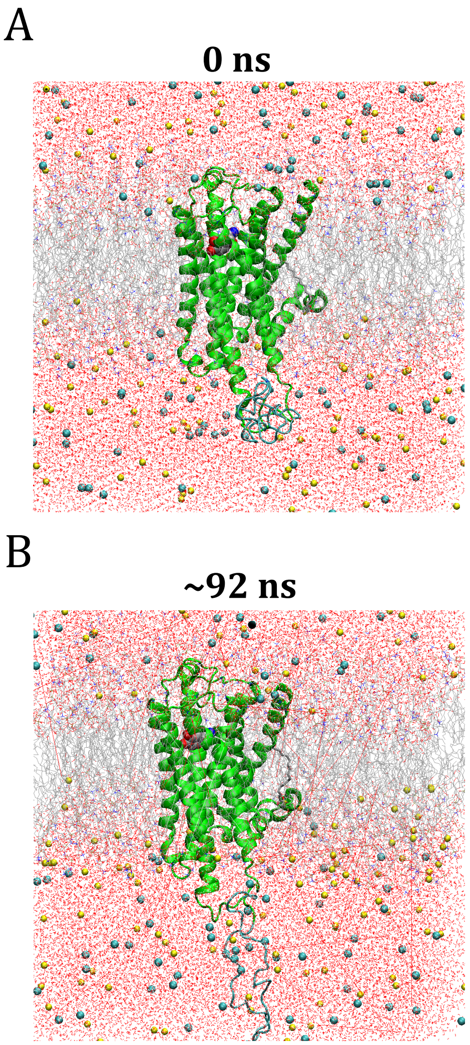


**<u>Figure S6</u>. Active human $\beta_1$AR structural model MD stability test.** All-atom MD simulation of the receptor bound to cationic norepinephrine embedded in a POPC/POPS lipid bilayer and hydrated by 0.15 M NaCl at 310 K and 1 atm, totaling ~187K atoms. NAMD 2.14 & 3.0 alpha (conducted on the Oracle cloud) was used to assess stability of the rebuilt model. Gradual protein harmonic restraints were applied for the first 42 ns. (**A**) Initial frame. **(B**) Final frame.
